# Polyunsaturated Fatty Acid - mediated Cellular Rejuvenation for Reversing Age-related Vision Decline

**DOI:** 10.1101/2024.07.01.601592

**Authors:** Fangyuan Gao, Emily Tom, Cezary Rydz, William Cho, Alexander V. Kolesnikov, Yutong Sha, Anastasios Papadam, Samantha Jafari, Andrew Joseph, Ava Ahanchi, Nika Balalaei Someh Saraei, David Lyon, Andrzej Foik, Qing Nie, Felix Grassmann, Vladimir J. Kefalov, Dorota Skowronska-Krawczyk

**Affiliations:** Gavin Herbert Eye Institute - Center for Translational Vision Research, Department of Ophthalmology, University of California Irvine, CA, 92697, USA; Department of Physiology and Biophysics, School of Medicine, University of California Irvine, CA; Department of Mathematics, University of California Irvine, CA; Department of Developmental and Cell Biology, University of California, Irvine, CA; The NSF-Simons Center for Multiscale Cell Fate Research, University of California Irvine, CA; Institute of Medical Sciences, University of Aberdeen, Aberdeen, UK; Department of Anatomy and Neurobiology, School of Medicine, University of California Irvine, CA; International Centre for Translational Eye Research, Institute of Physical Chemistry, Polish Academy of Sciences, Warsaw, Poland; Institute for Clinical Research and System Medicine, Health and Medical University, Potsdam, Germany

**Keywords:** PUFA, VLC-PUFA, ELOVL2, lipid supplementation, AMD

## Abstract

The retina is uniquely enriched in polyunsaturated fatty acids (PUFAs), which are primarily localized in cell membranes, where they govern membrane biophysical properties such as diffusion, permeability, domain formation, and curvature generation. During aging, alterations in lipid metabolism lead to reduced content of very long-chain PUFAs (VLC-PUFAs) in the retina, and this decline is associated with normal age-related visual decline and pathological age-related macular degeneration (AMD). *ELOVL2* (Elongation of very-long-chain fatty acids-like 2) encodes a transmembrane protein that produces precursors to docosahexaenoic acid (DHA) and VLC-PUFAs, and methylation level of its promoter is currently the best predictor of chronological age. Here, we show that mice lacking ELOVL2-specific enzymatic activity (*Elovl2^C234W^*) have impaired contrast sensitivity and slower rod response recovery following bright light exposure. Intravitreal supplementation with the direct product of ELOVL2, 24:5n-3, in aged animals significantly improved visual function and reduced accumulation of ApoE, HTRA1 and complement proteins in sub-RPE deposits. At the molecular level, the gene expression pattern observed in retinas supplemented with 24:5n-3 exhibited a partial rejuvenation profile, including decreased expression of aging-related genes and a transcriptomic signature of younger retina. Finally, we present the first human genetic data showing significant association of several variants in the human *ELOVL2* locus with the onset of intermediate AMD, underlying the translational significance of our findings. In sum, our study identifies novel therapeutic opportunities and defines ELOVL2 as a promising target for interventions aimed at preventing age-related vision loss.

## INTRODUCTION

The specific composition of lipids within membranes dictates their biophysical properties such as diffusion, permeability, domain formation, and curvature generation. Age-related changes in membrane lipid composition have been postulated to be one of the hallmarks of aging^1–3^. In the aged retina, polyunsaturated fatty acids (PUFAs), essential components of cellular membranes, show significantly decreased levels, and this decrease is further exacerbated in retinas affected by age-related macular degeneration (AMD)^4,5^. Retinal tissue is particularly enriched in long- and very long-chain PUFAs (LC-PUFA and VLC-PUFAs, respectively), which are integral components of photoreceptor disc membranes^6^, and their depletion or reduced levels have been postulated to be one of the hallmarks of AMD^7^.

Strategies aimed at preserving or replenishing VLC-PUFAs in the aging retina, such as dietary interventions with n-3 PUFAs and LC-PUFAs, are being investigated as potential approaches to maintain retinal health and function in older individuals ^8,9^. While some studies indicate improvement of vision following supplementation with docosahexaenoic fatty acid (DHA) or both DHA and eicosapentaenoic fatty acid (EPA)^10,11^, others, like Age-Related Eye Disease Study 2 (AREDS2), report no such correlation^12^, deeming DHA and EPA supplementation trials inconclusive. Interestingly, dietary and oral supplementation of animals with very high doses of VLC-PUFAs have shown promising results^10^. Another study has shown some improvement in visual performance in a dietary supplementation experiment with fish oil enriched in n-3 C24-28 VLC-PUFAs^11^. However, due to the high cost of synthesizing VLC-PUFAs and the limited accessibility to enriched fish oil, coupled with their relatively low efficacy, these methods currently lack practical applicability for preventing age-related decline in human vision.

The exact mechanisms underlying the decrease of VLC-PUFAs in aging and disease are not yet fully understood. The elongation and desaturation of essential fatty acids are tightly regulated and involve several enzymatic steps. The key enzyme involved in the elongation of LC- and VLC-PUFAs is ELOVL2 (Elongation of very long-chain fatty acids protein 2), which encodes an endoplasmic reticulum membrane-resident protein that produces precursors to DHA and VLC-PUFAs. *Elovl2* is primarily expressed in tissues with high metabolic demands, such as the liver, retina and brain^13,14^. Importantly, increasing methylation of the *ELOVL2* regulatory region has been shown to be one of the best biomarkers of chronological aging^15–17^. Recent work in our laboratory demonstrated that the expression and activity of ELOVL2 are closely linked to the aging process in the eye^14^.

In this study, we have identified and described critical structural, functional and molecular changes in the aging retina and correlated many of these phenotypes with the lack of ELOVL2 activity and disturbed lipid composition. We show that intravitreal injection of the direct product of ELOVL2 elongation, 24:5n-3, improves visual function, reduces severity of aging phenotypes, and decreases accumulation of deposits in aged mice. Notably, no other PUFA demonstrated such a profound effect. Finally, we present the first human genetic evidence of the impact of ELOVL2 activity or levels on risk of AMD. Our data underscores the importance of ELOVL2 activity in maintaining healthy vision and suggests a potential new therapy to reverse the symptoms of aging in the eye and prevent age-related eye diseases such as AMD.

## RESULTS

### Age-related vision decline is associated with decreased VLC-PUFA levels in the retina

To investigate changes in the lipid composition of aging retinas, we performed a series of lipidomic analyses on 3-, 6-, 12-, 18-, and 23-month-old dissected mouse tissues. For full lipidomic analysis, lipids were extracted using the Bligh-Dyer method^18^. For comprehensive fatty acids (FAs) analyses, FAs were released from lipids using the acid hydrolysis method and extracted using hexanes^19^. Complex lipids and FAs were analyzed by Liquid Chromatography Mass Spectrometry (LC-MS), as described in Materials and Methods. For FA analysis, the abundance of individual polyunsaturated fatty acids (PUFAs) was normalized to internal standard (FA 21:5) and tissue weight. Our data revealed a progressive decline in DHA (34% (*p*=0.0178) and 33% (*p*=0.0165) in 18-month-old and 23-month-old, respectively) and VLC-PUFAs, specifically 32:6 (31% in 18-month-old, *p*=0.0128), 34:6 (52% (*p*=0.0042) and 39% (*p*=0.0290) in 18-month-old and 23-month-old, respectively) and 36:6 (55% (*p*=0.0046) and 59% (*p*=0.0029) in 18-month-old and 23-month-old, respectively), in aged retinas compared to 3-month-old retinas, overall showing a significant decrease in levels of VLC-PUFAs in aging retina (**Fig. 1A)**.

**Fig. 1.**
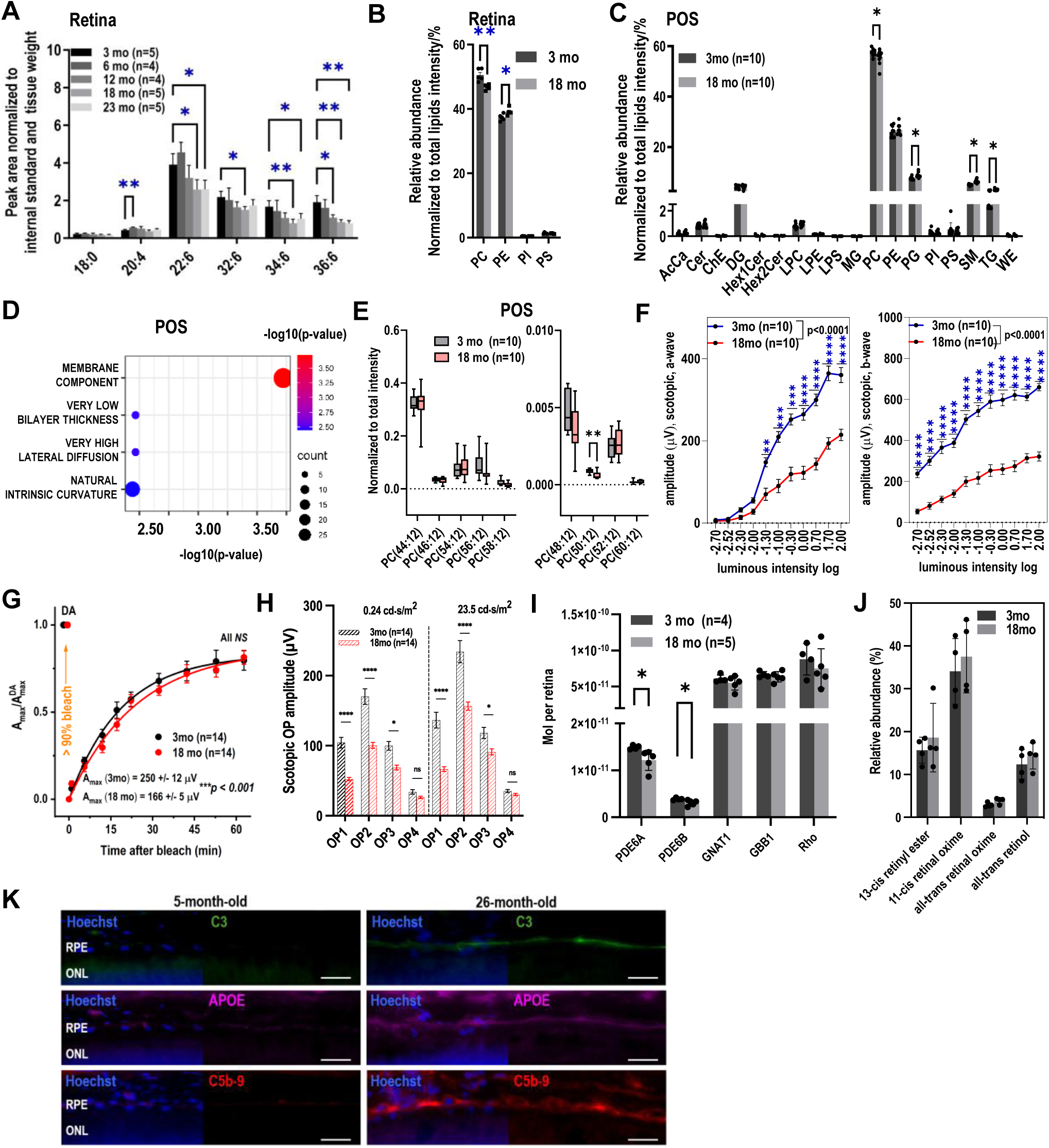
Age-related decreased VLC-PUFA levels in the retina is associated with vision decline. **(A)** Levels of 18:0, 20:4n-6, 22:6n-3 and VLC-PUFAs in retinas dependent on age (n=5, * = p<0.05, ** = p<0.01), **(B)** Levels of major phospholipid classes in 3-month-old and 18-month-old retinas (n=5, * = p<0.05, ** = p<0.01). **(C)** Quantification of lipid classes in aging photoreceptor outer segments (POS) (n=10, * = p<0.05). **(D**) Lipid ontology (LION) analysis of significantly changed lipids in aging POS showed severe alterations in membrane biophysical properties. (**E)** Down-regulated PC (48:12) and PC (50:12) in aging POS (n=10, * = p<0.05) **(F) S**cotopic electroretinogram (ERG) responses in aged (18-month-old) mice compared to young (3-month-old) mice (n=10, ** = p<0.01, *** = p<0.001, **** = p<0.0001). (**G**) Rate of rod-mediated dark adaptation recovery in young (3-month-old) and aged (17-month-old) mice (**H**) Oscillatory potential amplitudes of dark-adapted scotopic ERG in old (17-month-old) and young (3.5-month-old) mice (**I)** Absolute levels of PDE6A and PDE6B in aged (18-month-old) retinas quantified using a stable isotope–labeled (SIL) peptides-based method (n=5, * = p<0.05). (**J**) Relative abundance of retinoid species extracted from dark-adapted eyes in aged (18-month-old) mice compared to young (3-month-old) mice (n=4). (**K**) Immunostaining of C3, APOE and C5b-9, markers of AMD in young (5-month-old) and in old (26-month-old) retinas; scale bar 25um..

After conducting FA analysis, we explored the lipidomic changes in the aging mouse retina. The levels of extracted complex lipids were analyzed using a data-dependent acquisition (DDA) LC-MS method and lipid species were identified using LipidSearch software 4.2.21 (Thermo)^20^. Among the 528 lipids quantified in retinas, 15 lipids were significantly downregulated, and 53 lipids were upregulated in aging retinas (fold change > |1.5|, p-value <0.05) (**Fig. S1A**). Principal component analysis revealed clear clustering based on differences in lipid composition between 3- and 18-month-old retinas **(Fig. S1B)**. The major membrane lipid classes, phosphatidylcholine (PC) and phosphatidylethanolamine (PE), were significantly changed in 18-month-old retinas. Specifically, PC was significantly decreased (*p*=0.0039, n=5) and PE was significantly increased (*p*=0.0329, n=5) in aged retinas. **(Fig. 1B)**. Next, we isolated lipids from photoreceptor outer segments (POS) from 3- and 18-month-old mouse retinas and found a significant (5%) decrease in PC, and increase in phosphatidylglycerol (PG), sphingomyelin (SM) and triglycerides (TG) (18%, 26% and 25%, respectively) **(Fig. 1C)**. Based on the significantly changed lipids in POS membranes **(Fig. S1C**, fold change > |1.5|, p-value < 0.05), we performed lipid ontology (LION) enrichment analysis^21^, which revealed highly enriched changes in membrane components, as well as decreased bilayer thickness and high lateral diffusion in the membranes. (**Fig. 1D**). Since PC and PE are primarily present in plasma membranes, we conducted further analysis to assess the effects of aging on membrane composition. In particular, we focused on the levels of VLC-PUFAs incorporated in complex lipids and their impact on membranes in the aging retina. PC-VLC-PUFA levels were analyzed using Lipid Data Analyzer with a customized database. Our data revealed lower levels of these lipids; specifically, in POS, PC (50:12) and PE (48:12) decreased by 30% and 42%, respectively **(Fig. 1E, Fig. S1D)**.

To correlate age-related lipid changes with visual functions, a series of visual tests were performed on 3- and 18-month-old mice. First, quantitative optomotor response (OMR) analysis was performed to measure contrast sensitivity in scotopic conditions to mimic nighttime light levels. Unsurprisingly, 18-month-old mice had significantly lower contrast sensitivity in scotopic conditions compared to 3-month-old mice (**Fig. S1E)**. Using electroretinography (ERG), we demonstrated a significant decrease in both scotopic a- and b-waves (**Fig. 1F**). In full-field ERG stimulation, scotopic a-wave was reduced by ∼ 34% in 18-month-old mice (250 ± 12 µV, n = 14) as compared to that in 3-month-old animals (166 ± 5 µV, n = 14, \*\*\**P* < 0.001). Photopic green/UV a- and b-waves were also reduced in 18-month-old mice compared to 3-month-old mice (**Fig. S1F, S1G**).We then investigated whether the dark adaptation of rod photoreceptors is suppressed in aged mice. After bleaching ∼90% of their visual pigment, rods in both groups gradually recovered their photoresponses over the following 60-min period in the dark. The recovery of the averaged *A*_max_ in young rods could be described by a single exponential function with a time constant of 18.9 ± 0.7 min, and its level by 60 min after the bleach was ∼ 80 ± 6% of the pre-bleach value (**Fig. S1H**). Although rods in aged mice also demonstrated robust recovery of their photovoltage after the bleach, its rate (24.3 ± 2.7 min) was slightly decreased (by ∼ 1.3 times, p<0.001) as compared to that in young mouse rods. Yet, the maximal response amplitude 60 min post-bleach reached the same relative level (∼ 81 ± 4%) as in young control rods (**Fig. 1G**). We also compared the oscillatory potentials in young and old mice and found a consistent significant reduction in all four oscillatory potential peaks in both dim flash and bright flash scotopic responses (**Fig. S1G, Fig. 1H**). However, the recovery of rod-driven ERG a-wave sensitivity (*S*_f_) following the same bleach was not compromised in older animals (**Fig. S1I**), suggesting the normal operation of their visual cycle. Taken together, our results show that aging affected visual function through decreased contrast sensitivity and decreased photoreceptor function.

To address whether the decrease in visual function in aged animals could be attributed to alterations in the phototransduction cascade or visual cycle, we performed two assays. First, we quantified levels of proteins involved in signal transduction in POS, including Rho, PDE6A, PDE6B, GNAT1 and GBB1, using a stable isotope-labeled (SIL) peptides-based absolute quantification method. To compare the levels of PDE6 and transducin proteins in 3- and 18-month-old retinas, the optimal amount of SIL peptides was spiked into the protein samples, trypsinized and analyzed by LC-MS/MS. The amount of each protein was calculated using the endogenous/SIL peptides peak area ratio and the quantification results were normalized to Rho and displayed as the mean ± standard error of the mean. Absolute quantification of these proteins showed slight decreases in PDE6A and PDE6B in 18-month-old retinas (**Fig. 1I**); however, relative amounts of all measured proteins (PDE6A, PDE6B, GNAT1 and GBB1) normalized to levels of Rhodopsin (RHO) remained unchanged (**Fig. S1J**). This suggests that the ratio of proteins involved in the phototransduction signaling pathway was not affected in aged retina. To investigate the impact of aging on overall visual cycle capacity, we then extracted retinoids from dark-adapted eyes following a previously published protocol^22^. Consistent with the normal recovery of sensitivity (*S*_f_) following a bleach, relative amounts of retinoid species showed no significant differences between 3- and 18-month-old eyes **(Fig. 1J)**.

Finally, to evaluate age-related changes in levels of proteins associated with retinal pathologies, we performed immunofluorescence staining on retinal cross-sections from young (5-month-old) and aged (26-month-old) mice. We used antibodies targeting Complement factor 3 (C3), Apolipoprotein E (ApoE) and subunit of the membrane attack complex (MAC) in the complement pathway C5b-9, all of which are correlated with increased risk of AMD. Our data demonstrated higher levels of these proteins in the RPE cell layer of 26-month-animals when compared to 5-month-old retina sections (**Fig. 1K**). Additionally, immunofluorescence staining with rod bipolar- and Müller cell-specific antibodies (protein kinase C alpha, PKCa and glutamine synthetase, GS, respectively) showed age-related morphological changes in the outer plexiform layer including overgrowth of dendrites into the outer nuclear layer (**Figure S1K, L**).

### Age-related changes in cone Elovl2 expression

*ELOVL2* encodes a key enzyme in the biosynthesis of long-chain fatty acids **(Fig. 2A**). Specifically, it elongates 22:5n-3 to 24:5n-3, which is then elongated to VLC-PUFAs by other elongases including ELOVL4 or further processed in peroxisomes to produce 22:6n-3, DHA. Our previous data indicated higher methylation levels in the *Elovl2* regulatory region in aged retina^14^. Here, we analyzed methylation levels at each CpG site in the *Elovl2* promoter that has been shown to undergo methylation changes with age in human blood^15^ **(Fig. 2B)**. To achieve this, we isolated DNA from young (<12-month-old) and old (24-month-old) mouse retinas and subjected it to bisulfite conversion, a process which involves treatment with sodium bisulfite to distinguish methylated from unmethylated cytosines. Then, using both converted and non-converted DNA, we amplified regions of interest with primers insensitive for methylation. PCR products were then sequenced using Sanger sequencing and relative levels of methylated versus non-methylated cytosines were calculated by comparing the area under the sequencing peak in both DNA samples. Our data revealed a group of five CpGs carrying significantly higher methylation levels in older retinas, while the other four, the closest to the transcription start site, were not changed. Notably, the CpG sites that exhibited significantly higher methylation in mice are direct homologs of CpGs methylated in human aging blood^23^.

**Fig. 2.**
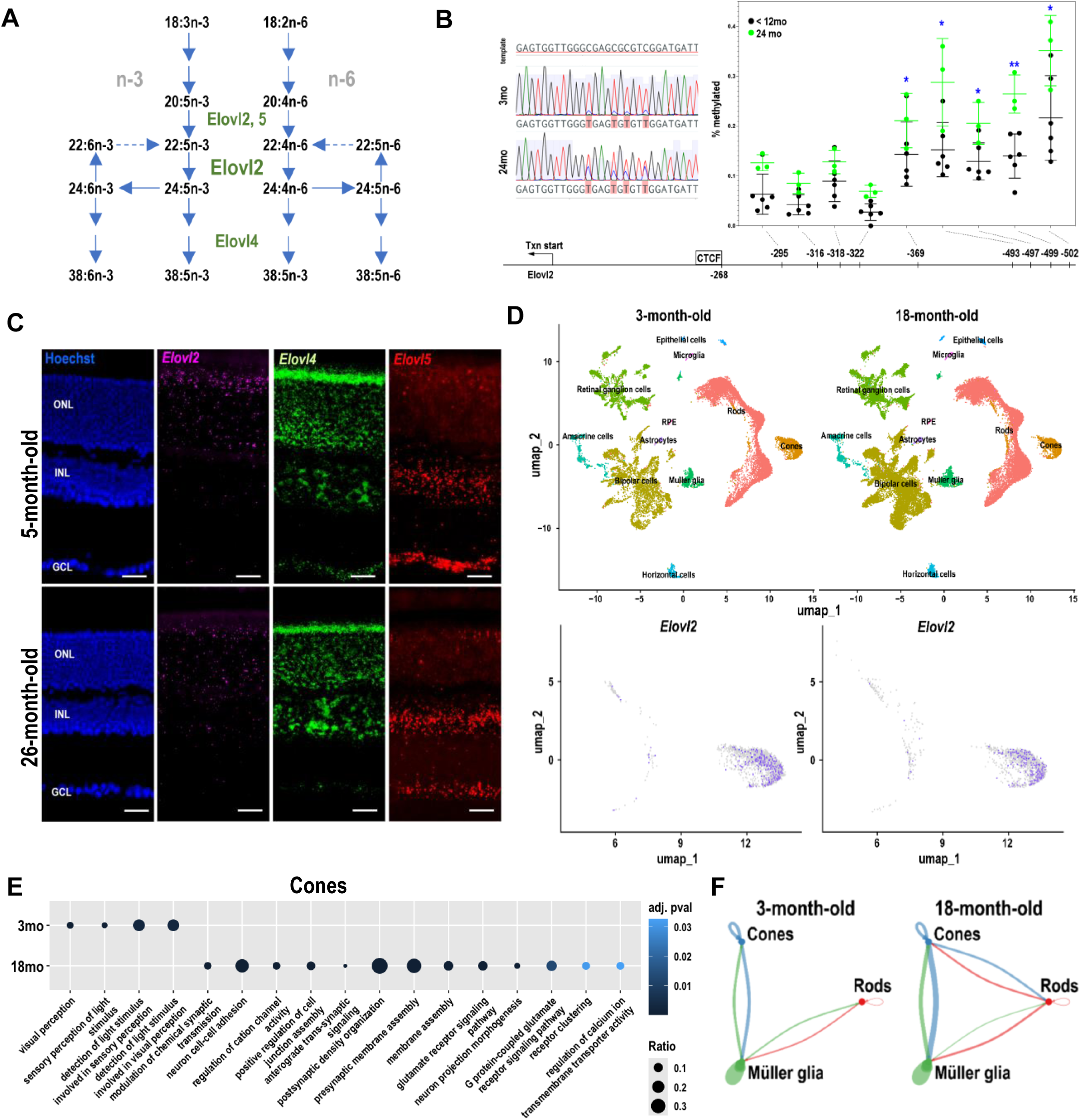
Age-related decrease in *Elovl2* expression. (**A**) VLC-PUFA elongation pathways. VLC-PUFAs are produced from essential FAs elongation by enzymes including Elovl2, 4, 5. (**B**) *Elovl2* promoter is increasingly methylated with age in mouse retina. (**C**) Images of mouse retina sections from young—5mo (top panels) and old—26mo (bottom panels) animals stained with RNAscope probes designed for *Elovl2*, *Elovl4* and *Elovl5*, counterstained with Hoechst. (**D**) snRNA-seq demonstrated decreased expression of *Elovl2* in cone photoreceptors in old (18-month-old) mice compared to young (3-month-old) mice..(**E**) GSEA analysis of pathways enriched in cones from young (3-month-old) and old (18-month-old) mice. (**F**) Visualization of interaction strength across young and old retinas. Each plot represents interactions among the cell types. The thickness of the lines between the cells indicates the interaction strength, with thicker lines representing stronger interactions.

To visualize cell type- and age-specific expression of key enzymes in the PUFA elongation pathway, we performed RNA *in situ* hybridization on retinal cross-sections from adult (5-month-old) and advanced-aged (26-month-old) mice using RNAscope probes targeting *Elovl2*, *Elovl4*, and *Elovl5* **(Fig. 2C)**. *Elovl2* expression was observed in the photoreceptor layer, whereas *Elovl4* was expressed in all retinal layers, and *Elovl5* was expressed in inner nuclear and ganglion cell layers. While *Elovl4* and *Elovl5* expression remained relatively unchanged with age, *Elovl2* expression decreased in 26-month-old animals compared to 5-month-old animals, as previously reported^14^.

To identify the specific cell types expressing *Elovl2* and other PUFA elongation enzymes, we performed single-nucleus RNA sequencing (snRNAseq) on retinas isolated from young (3-month-old) and old (18-month-old) mice. After normalizing mean expression levels to TBP, *Elovl2* expression was significantly decreased in aged retinas (0.080 vs. 0.100 in young retinas, p-value = 3.59E-6). As previously observed in human retinas^24^, expression of *Elovl2* was highest in cones and barely detectable in other cell types (**Fig. 2D**). Our data is in agreement with data obtained by the CZI CELL×GENE Discover program^25^ from mouse and human retinas (**Fig S2A**), confirming *Elovl2* enrichment in cones. In addition, we investigated the cell type-specific expression of other enzymes from the PUFA elongation pathway and discovered that only *Elovl2* exhibited cone-specific expression, while other enzymes were less restricted. Based on these findings, we subsequently investigated the pathways that undergo alterations in cone photoreceptors during aging. Differentially expressed genes (DEGs) (log fold change > 0.1, padj < 0.05) were identified from young (3-month-old) and aged (18-month-old) cones, which were then used in GO functional enrichment analysis **(Fig. 2E)**. Biological pathways that were enriched in young cones were related to visual perception and light stimulus detection. In aged cones, pathways that were enriched were involved in synaptic transmission, ion channels, and membrane assembly.

To gain deeper insights into the evolution of intercellular interactions in the aging retina, we employed the computational package CellChat^26^, focusing specifically on interactions between cones, rods and Müller glia (**Fig. 2F, S2B, S2C**). Using our snRNA-seq data, we first evaluated the overall inferred number and strength of interactions within and between the cell types. In the young retinas, the strongest interactions were detected between Müller glia and cones. These interactions were further increased in aged tissue. Similarly, signaling between rod and Müller glia, increased in aged retina. Interestingly, cone-rod interactions were not detected in young retinas, while several new significant interactions were detected in 18-month-old tissues. The analysis of specific ligand-receptor pairs within these signaling interactions revealed the emergence of interactions between neuronal adhesion molecules (*Negr1 - Negr1*) and metabotropic glutamate receptors (*Glu - Grm8*) between rods and cones, specifically in aged retinas. In addition, several interactions between metabotropic glutamate receptors (*Glu - Grm8, Glu - Grm7*) and ionotropic glutamate receptors (*Glu - Gria4*) emerged within Müller glia in aged retinas **(Fig. S2B, S2C)**. In addition, our analysis showed new Cyclosporin A and CD147 (*Ppia* - *Bsg*) interactions between Müller and photoreceptor cells, which are recognized as an infection sensor and an inflammation initiation system^27^ (**Fig. S2B**).

### Lack of ELOVL2 activity accelerates changes in age-related phenotypes in the retina

In our previous work, we generated Elovl2-mutant mice (*Elovl2^C234W^*) with loss of ELOVL2 enzymatic activity due to impaired substrate binding ability^14^. Here, we performed detailed analysis of lipid composition, including total fatty acids and global lipidomic analysis on retinas from 18-month-old *Elovl2^C234W^* mice. The quantification revealed that levels of PUFAs synthesized from ELOVL2 products, including 22:6, 24:6, 32:6, 34:6 and 36:6 were significantly lower in 18-month-old *Elovl2^C234W^* retinas than in age-matched wildtype retinas (**Fig. 3A)**. Untargeted lipidomic analysis of retina was also performed to compare lipid species from 18-month-old *Elovl2^C234W^* and wild-type retinas. Of the 595 lipids identified, 28 were significantly downregulated and 84 were significantly upregulated in *Elovl2^C234W^* retinas (fold change > |1.5|, p-value < 0.05) **(Fig. S3A)**. In addition, *Elovl2^C234W^* tissue displayed a significant decrease in PC and increase in PE levels (**Fig. 3B)**, similar to the changes detected in aging retinas. Lipid ontology (LION) enrichment analysis based on the significantly changed lipids suggested highly enriched changes in plasma membrane components, as well as decreased bilayer thickness and high lateral diffusion, which were also similar to the changes in aging POS **(Fig. 3C)**. Therefore, we further analyzed VLC-PCs and the results showed decreased PC(46:12), PC (54:12), PC (56:12) **(Fig. 3D)**, which contributed to decreased total level of VLC-PCs in 18-month-old *Elovl2^C234W^* retina **(Fig. S3B)**. Analysis of free fatty acids (FFA) showed 45% (*p* = 0.048), 70% (*p* = 0.002), 92% (*p* = 0.002), 62% (*p* = 0.008), 57% (*p* = 0.008) and 32% (*p* = 0.128) decrease of 22:6, 24:6, 26:6, 32:6, 34:6 and 36:6 in 18-month-old *Elovl2^C234W^* retinas compared to age-matched wildtype retinas **(Fig. S3C)**.

**Fig. 3.**
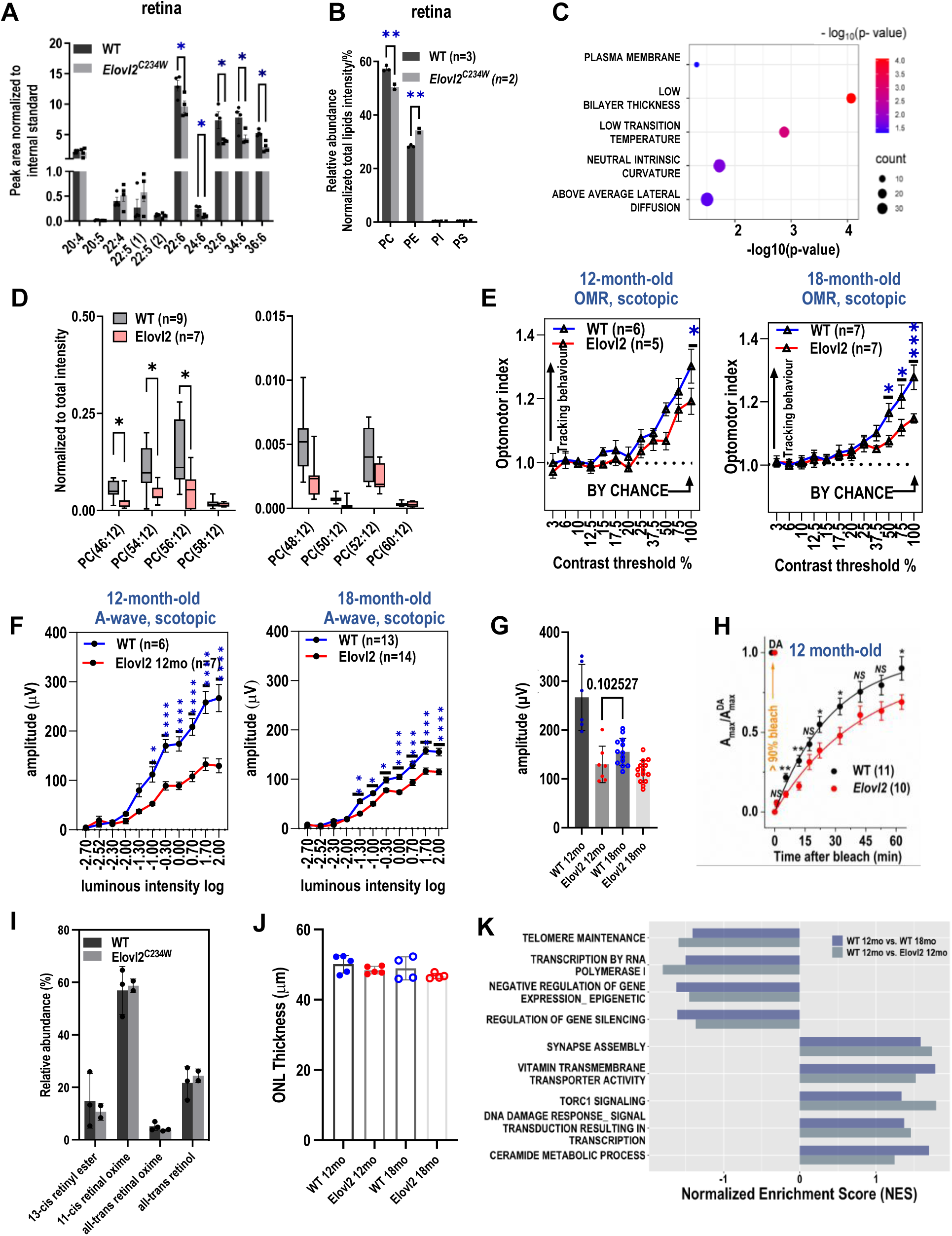
Disturbed lipid composition in Elovl2^C234W^ mouse retinas is correlated with vision loss. (**A**) Total fatty acid products of ELOVL2 elongation, DHA (22:6) and VLC-PUFAs (24:6, 32:6, 34:6 and 36:6) were decreased in retinas of Elovl2^C234W^ mice compared to age-matched wildtype mice (n=4, * = p<0.05). (**B**) Changes in major phospholipid classes in Elovl2^C234W^ retinas compared to age-matched wildtype retinas resemble changes seen in aged retinas compared to young retinas (** = p<0.01). (**C**) Lipid ontology (LION) analysis of significantly changed lipids in Elovl2^C234W^ retinas showed severe alterations in membrane biophysical properties. (**D)** VLC-PC species in Elovl2^C234W^ retinas (* = p<0.05). (**E**) Decreased contrast sensitivity, (**F**) scotopic electroretinogram (ERG) responses and (**H**) delayed rod-mediated dark adaptation of Elovl2^C234W^ mice compared to age-matched wildtype mice (* = p<0.05, ** = p<0.01, *** = p<0.001, **** = p<0.0001). (**G**) Maximum ERG a-wave amplitudes of 12-month-old Elovl2^C234W^ retinas were similar to those of 18-month-old wild-type retinas. (**I**) No difference in outer nuclear layer (ONL) thickness between 12-month-old wildtype and Elovl2^C234W^ retinas as measure by OCT (n=5/group), and 18-month-old Elovl2^C234W^ and wildtype retinas (n=4/group) (**J**) Relative abundance of retinoid species extracted from dark-adapted eyes remained unchanged in 18-month-old Elovl2^C234W^ (n=2) and wildtype (n=3) retinas. (**K**) Comparison of GSEA Normalized Enrichment Scores (NES) of selected GO terms in 12-month-old Elovl2C234W vs. 12-month-old wildtype retinas with the NES of GO terms in 18-month-old vs. 12-month-old retinas.

Next, we compared visual function in 12- and 18-month-old *Elovl2^C234W^* mice. OMR analysis showed significantly lower scotopic contrast sensitivity in *Elovl2^C234W^* animals compared to age-matched wildtype animals at both ages **(Fig. 3E)**. Scotopic a- and b-wave ERG responses were also reduced at both ages compared to wildtype controls **(Fig. 3F, Fig. S3D)**. Notably, the maximum ERG a- and b-wave amplitudes of 12-month-old Elovl2^C234W^ (120 μV) retinas were decreased to levels comparable to those of 18-month-old wild-type (150 μV, p = 0.103) retinas (**Fig. 3G, Fig. S3D**). There were no significant differences in a- and b-wave photopic and c-wave ERG amplitudes between wildtype and *Elovl2^C234W^* animals (**Fig. S3E, F**).

The rod dark adaptation in ELOVL2-deficient mice was then evaluated after a nearly complete rhodopsin bleach with green light. In control mice, the averaged scotopic a-wave maximal response, *A*_max,_ recovered with the time constant of 28.6 ± 2.6 min, and its level by 60 min after the bleach was ∼ 90 ± 7% of the pre-bleach value (**Fig. 3H**, black symbols). In contrast, the recovery of the scotopic a-wave response in *Elovl2^C234W^* mice was significantly suppressed compared to controls. Consistent with this, the average rate of rod *A*_max_ recovery in *Elovl2^C234W^* mice (42.7 ± 7.3 min) was ∼ 1.5 times slower than in control animals (28.6 ± 2.5 min, *p*=0.000008), and reached only ∼ 69 ± 5% of its prebleached level by the end of 60-min recordings (**Fig. 3H**, red symbols). The dark-adapted a-wave photosensitivities (*S*_f_) were also lower (by ∼ 16%) in the same group of mice lacking ELOVL2 (1.50 ± 0.05 m^2^ cd^-1^ s^-1^ vs. 1.79 ± 0.08 m^2^ cd^-1^ s^-1^ in controls, \**P* < 0.05). However, the recovery of rod-driven *S*_f_ following the same bleach was not suppressed in ELOVL2-deficient animals **(Fig. S3G)**, suggesting the normal recycling of the visual chromophore in the mutant mice.

To investigate further whether loss of ELOVL2 function affects overall visual cycle capacity, we extracted retinoids from dark-adapted eyes from 18-month-old *Elovl2^C234W^* and age-matched wildtype animals **(Fig. 3I)**. Consistent with the normal recovery of sensitivity *S*_f_ following a bleach, quantification of the relative amounts of retinoid species revealed no significant difference between *Elovl2^C234W^* and wildtype animals. To determine whether the reduction in ERG amplitude in *Elovl2^C234W^* retinas could be due to cell loss or retinal degeneration, we measured outer nuclear layer (photoreceptor) thickness by optical coherence tomography (OCT). We found no statistically significant difference by 2-way ANOVA in ONL thickness between 12-month-old wildtype and *Elovl2^C234W^* (n=5), and 18-month-old wildtype and *Elovl2^C234W^* (n=4) retinas **(Fig. 3J)**. Therefore, the reduction in photoreceptor function in 12-month-old *Elovl2^C234W^* retinas preceded any significant cell loss or retinal degeneration.

Given that *Elovl2^C234W^* animals exhibited accelerated aging phenotypes in our data, we performed bulk RNA sequencing on 12-month-old wildtype and *Elovl2^C234W^* retinas. We then compared this data to the transcriptomes of 12-month-old and 18-month-old wildtype retinas to identify potential transcriptomic parallels in retinal changes associated with the absence of the ELOVL2 enzyme and in aging. Gene set enrichment analysis (GSEA) was performed for differentially expressed genes (fold change > |1|, padj < 0.05) using Gene Ontology Biological Processes, Cellular Component, and Molecular Function gene sets. Pathways that were downregulated in both aging retinas and retinas lacking ELOVL2 included telomere maintenance, transcription by RNA polymerase I, negative regulation of gene expression (epigenetic), and regulation of gene silencing **(Fig. 3K)**. On the other hand, pathways such as synapse assembly, vitamin transmembrane transporter activity, TORC1 signaling, DNA damage response-signal transduction resulting in transcription, and ceramide metabolic process were upregulated in both 12-month-old *Elovl2^C234W^* and 18-month-old wildtype retinas when compared to 12-month-old wildtype retinas **(Fig. 3K, S3H)**.

### Rescue of age-related vision decline by intravitreal fatty acid injection

We hypothesized that the lack of ELOVL2 product, 24:5n-3, in the aging retina is one of the main culprits of age-related vision decline and that supplementation with this fatty acid may improve vision in aged animals. To test this hypothesis, we pursued an intravitreal supplementation strategy, which, in contrast to oral gavage, enabled us to control precisely the amount of lipid administered in the eye. To assess the retinal toxicity of 24:5n-3, intravitreal injections of 24:5n-3 were administered unilaterally to 3-month-old mice, and ERG responses were measured 2 days and 5 days post-injection (**Fig. S4A**). Compared to vehicle-injected eyes, 24:5n-3-injected eyes did not show any significant change in scotopic a- or b-wave ERG amplitudes on day 2 (**Fig. S4B**). Additionally, no significant changes were observed in either scotopic or photopic responses on day 5 (**Fig. S4C**). In summary, intravitreal injection of 24:5n-3 in 3-month-old mice did not demonstrate retinal toxicity and did not alter rod- or cone-driven retinal responses.

To investigate the effect of 24:5n-3 supplementation on visual function in aged mice, we administered an intravitreal injection of 24:5n-3 into one eye of each 18-month-old animal, while the contralateral eye received an injection of a vehicle as a control. After 5 days, a multimodal approach was used to test impact on visual function, and the retinas were subsequently collected for analysis of fatty acids and lipid composition, and investigation of molecular changes **(Fig. 4A)**. First, we tested different doses of 24:5n-3 to determine the optimal levels that could improve vision in 18-month-old mice. Interestingly, we observed that 24:5n-3-treated eyes showed statistically significant functional improvement in both photopic and scotopic a- and b-wave ERG responses with an injection dose of 0.36 nmol (**Fig. 4B**), and no significant changes were observed when other doses were used (**Fig. S4E**). Then, to test whether PUFAs up- or downstream of 24:5n-3, such as 20:5n-3, 22:6n-3 and 32:6n-3, have similar effects, we replicated the procedure using the optimal dose (0.36 nmol) and administered intravitreal injections of 20:5n-3, 22:6n-3 and 32:6n-3, followed by functional analysis **(Fig. 4B, S4F)**. Strikingly, only eyes that received intravitreal supplementation of 24:5n-3 exhibited statistically significant functional improvement in both photopic and scotopic a- and b-wave ERG responses when assessed 5 days post-injection. In contrast, injection of 22:6n-3 and 20:5n-3 resulted in no functional improvement in photopic and scotopic ERG responses (**Fig. 4B, S4F**). A marginal improvement in photopic green and blue b-wave responses, but not in scotopic responses, was also observed in eyes treated with intravitreal administration of 32:6n-3 (**Fig. S4F**).

**Fig. 4.**
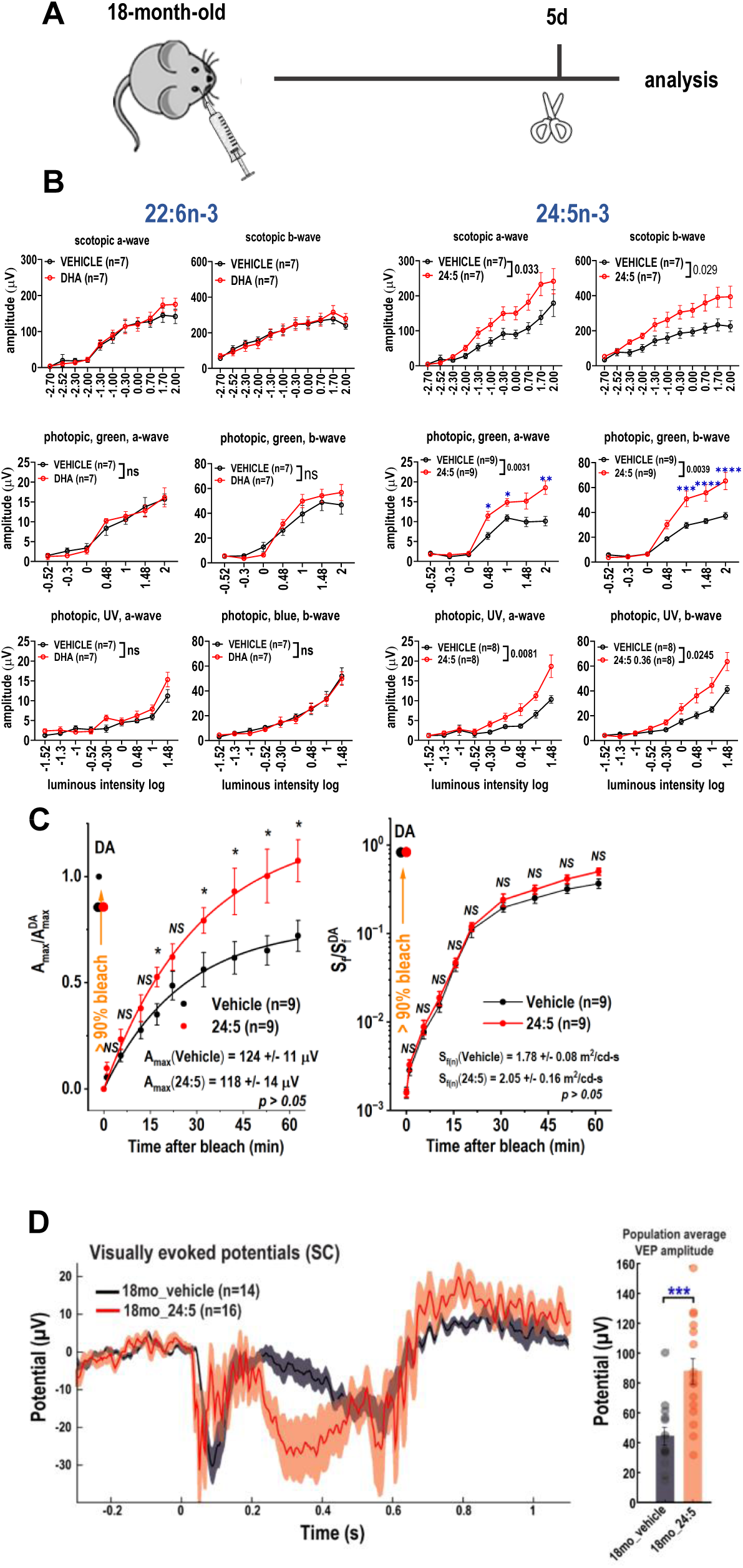
Intravitreal supplementation of 24:5n-3 in aged mice rescues visual function. (**A**) Schematic of intravitreal supplementation of 24:5n-3 in 18-month-old mice. (**B**) Improvement of scotopic and photopic a- and b-wave electroretinogram (ERG) responses, **(C)** faster rod-mediated dark adaptation recovery, and (**D**) improved visual-evoked potentials (VEP) 5 days following intravitreal supplementation of 0.36 nmol 24:5n-3 (*=p<0.05, **=p<0.01; ***=p<0.005; ****=p<0.001), but not of 0.36nmol 22:6n-3 (DHA).

The effect of intraocular 24:5n-3 supplementation on rod dark adaptation was assessed by *in vivo* ERG. In 17-month-old mice, this treatment improved the final post bleach recovery fraction of rod *A*_max_ response by ∼ 30% (**Fig. 4C**, left), as compared to that in vehicle-injected contralateral eyes, without affecting their sensitivity recovery **(Fig. 4C**, right). In contrast, a similar administration of 24:5n-3 did not affect the recovery of either of the two parameters in 3.5-month-old animals (**Fig. S4D**), thus indicating the specificity of the therapeutic effect of the lipid to the aged mice.

Finally, we evaluated the effects of 24:5n-3 supplementation on the health of the visual extrageniculate (via the Superior colliculus; SC) pathway by recording visually evoked potentials (VEP). Our recordings in the SC showed a significant increase in VEP peak-to-peak amplitude in 18-month-old animals supplemented with 24:5n-3 compared to the vehicle-treated eyes **(Fig. 4D)**. Specifically, the VEP amplitude increased from 44.15 ± 5.95 µV in 18-month-old animals injected with a vehicle (n=4 animals; n=14 recording sites) to 86.67 ± 8.61 µV in supplemented eyes (n=4; n=16; p-value = 0.00082).

### Mechanism of vision improvement upon intravitreal 24:5n-3 supplementation

To gain insight into the potential mechanism underlying the improvement in vision following 24:5n-3 supplementation, we first investigated whether the lipid supplementation could cause any changes in lipid composition in the retina. Tissues were collected 5 days after the vehicle or lipid injection, and lipids were extracted using the Bligh and Dyer method. We analyzed the levels of all possible VLC-PUFA-containing phospholipids from isolated POS and noted that all were elevated in the 24:5n-3 injected group. Among them, VLC-PUFA incorporated PE, PC, LPC, PS, and plasmalogen PE increased significantly by 25% (*p*=0.035), 26% (*p*=0.070), 20% (*p*=0.017), 22% (*p*=0.025), 40% (*p*=0.007), in the 24:5n-3-supplemented group, respectively (**Fig. 5A**). Overall, the level of phospholipids containing VLC-PUFAs was significantly elevated by ∼25% (*p*=0.033) in the retinas receiving fatty acid supplementation. Analysis of the FA levels, however, indicated no change including no increase in total levels of VLC-PUFAs following supplementation (**Fig. S5A, B**).

**Fig. 5.**
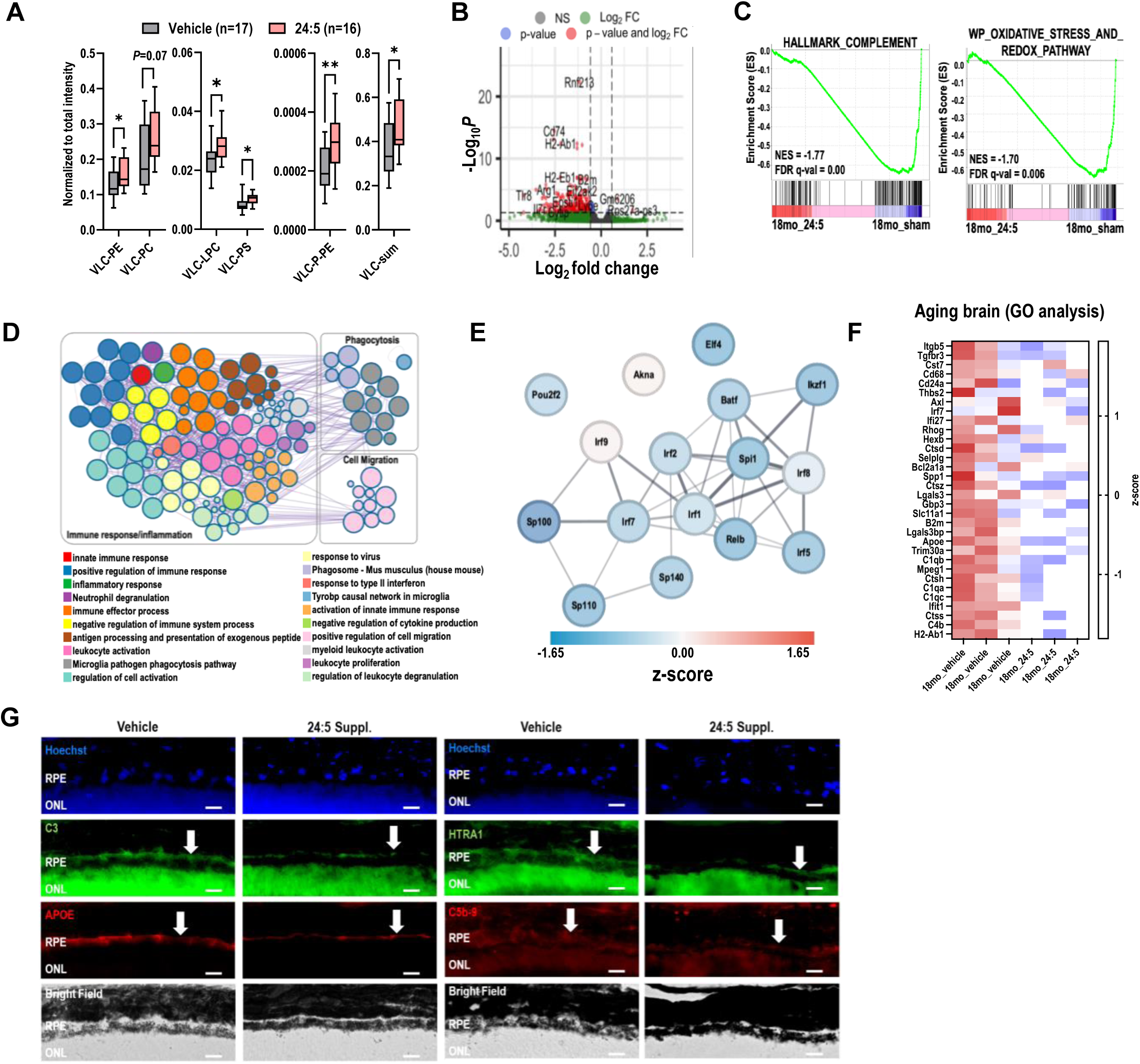
Reversal of molecular aging phenotypes in 18-month-old 24:5n-3-supplemented eyes. **(A)** Increased levels of possible classes of VLC-PUFA-incorporated phospholipids in isolated photoreceptor outer segments (POS) following intravitreal supplementation of 24:5n-3 (* = p<0.05, ** = p<0.01). **(B)** Nearly all differentially expressed genes (DEGs) (FC > |1.5|, pval<0.05) in 18-month-old 24:5(n-3)-supplemented retinas were downregulated compared to vehicle-injected retinas (n=3). **(C)** Gene Set Enrichment Analysis (GSEA) enrichment plots revealed downregulation of complement (NES=-1.77, FDR qval=0.00) and oxidative stress and redox (NES=-1.70, FDR qval=0.006) pathways following 24:5(n-3) supplementation (n=3). **(D)** Metascape analysis demonstrated downregulation of immune response, inflammation, microglial phagocytosis and cell migration pathways following 24:5n-3 supplementation (n=3). **(E)** STRING functional protein-protein interaction network of top 20 transcription factors involved in gene regulation after 24:5n-3 supplementation. **(F)** Downregulation of genes in the aging cerebellum gene set following 24:5n-3 supplementation (n=3). **(G**) Immunofluorescence staining shows lower expression of complement C3, APOE, HTRA1 and C5b-9 proteins in RPE cell layer in retinal cross-sections from 24:5n-3 supplemented eyes.

To understand the molecular changes occurring in the retina after fatty acid supplementation, we performed bulk RNAseq analysis on vehicle- and 24:5n-3-injected 18-month-old retinas isolated five days following intravitreal injection. The changes in the transcriptome were identified using Bioconductor package DESeq2^28^. We identified 234 significantly changed genes (fold change > |1.5|, p-adj < 0.05). Interestingly, almost all differentially expressed genes (DEGs) were downregulated in the 24:5n-3-supplemented eyes **(Fig. 5B)**. Gene set enrichment analysis (GSEA) indicated significant downregulation of complement (NES = -1.77, FDR q-value = 0.00) and oxidative stress and redox pathways (NES = -1.70, FDR q-value = 0.006) in 24:5n-3-supplemented retinas **(Fig. 5C)**. The visualization of enriched terms as a network using Metascape^29^ demonstrated downregulation of pathways involved in immune response, inflammation, microglial phagocytosis, and cell migration **(Fig. 5D, S5C).**

To further identify transcription factors (TFs) responsible for the observed gene expression changes following 24:5n-3 supplementation, we used ChIP-X Enrichment Analysis 3 (ChEA3), a tool that predicts TFs associated with DEGs based on TF-target gene set libraries curated from human, mouse and rat^30^. We used the Mean Rank output, which averaged integrated ranks across libraries, for downstream analysis and created global TF co-expression networks. Our RNA sequencing data demonstrated downregulation of *Sp100, Lyl1, Sp140, Irf5, Stat6, Fli1, Ikzf1, Spi1, Relb, Elf4, Sp110* and upregulation of *Trafd1* **(Fig. S5D)**. To visualize protein-protein interactions among these associated TFs, we created a functional interaction network of the top 20 TFs using STRING (v12.0)^31^ **(Fig. 5E)**. Low expressing TFs, as defined by normalized read count (DESeq2 baseMean) < 10, were removed, and the difference in mean Z-scores after 24:5n-3 supplementation of the remaining 16 TFs were calculated. Our data demonstrated striking downregulation of nearly all top TFs, including several interferon regulatory factors (IRFs), *Irf1, Irf2, Irf5, Irf7, Irf8,* responsible for activating immune response^32^. Interestingly, expression of genes related to rod and cone photoreceptor-specific ion channels remained unchanged in retinas that received 24:5n-3 supplementation compared to vehicle-injected retinas, suggesting unaffected phototransduction in 24:5n-3 supplemented eyes **(Fig. S5E)**.

Further analysis demonstrated significant downregulation of genes in the “aging cerebellum” gene set^33^ (NES = -1.82, FDR = 5.83E-4) in 18-month-old retinas after supplementation with 24:5n-3 compared to vehicle, including significant downregulation of apolipoprotein E (*Apoe)* transcript (log2FC = -0.79, p-adj = 0.001) **(Fig. 5F)**. To corroborate our RNA sequencing data showing decreased expression of *Apoe* and genes related to the complement cascade after supplementation, we performed immunohistochemical analysis on retinal cross-sections **(Fig. 5G)**. First, we observed a marked reduction of complement component C3, a key inflammatory protein activated in AMD^34^, in the RPE of 24:5n-3-supplemented eyes. We also observed decreased accumulation of APOE, a major component of age-related sub-RPE drusenoid deposits^35^. Levels of HTRA1 protein, a serine protease secreted by RPE that is increased in AMD^36^, was found to be reduced following treatment with 24:5n-3 treatment. Lastly, we observed decreased levels of the C5b-9, subunit of the membrane attack complex, the terminal complex of the complement cascade^37^, indicating decreased complement activation in 24:5n-3-supplemented eyes.

### Genetic connection between aging and AMD

Our studies revealed the crucial role of ELOVL2 in preserving retinal health in pre-degenerative stages. Therefore, we investigated the association between genetic variants within the *ELOVL2* (chr6: 10980992 – 11044624) gene and the age of onset of intermediate age-related macular degeneration (AMD) using available human population data. The association between age of onset and genetic variants within the *ELOVL2* gene body was assessed in two cohorts: unrelated, European individuals with intermediate AMD from the International AMD Genomics Consortium (IAMGDC, n=2,407, mean age at diagnosis: 74.1 years)^38^, as well as incident AMD cases from the UK Biobank (n= 1,309, mean age of onset: 62.8 years). In the IAMDGC cohort, we used linear regression with age of onset as the outcome and genotype as the exposure, adjusted for study, sex and the first five principal components of ancestry. In the UK Biobank, the analyses were additionally adjusted by smoking status and BMI.

The strongest association was observed for rs911196 in the fifth intron of *ELOVL2*. The minor G allele (allele frequency 25% in Europeans) resulted in 4.7 months (95% Confidence Interval: 2.1 months – 7.3 months) earlier onset for intermediate AMD in our analyses (P=0.0003, **Fig. 6**). This effect was seen in both cohorts (IAMDGC: 5.7 months earlier onset, UK Biobank: 4.5 months earlier). The G allele of the correlated variant rs9468304 in the first intron also showed a significant association, however the effect size was weaker (31% allele frequency, 4.32 months earlier onset, P= 0.0009). None of the remaining variants had a statistically significant correlation (P>0.05).

**Fig. 6.**
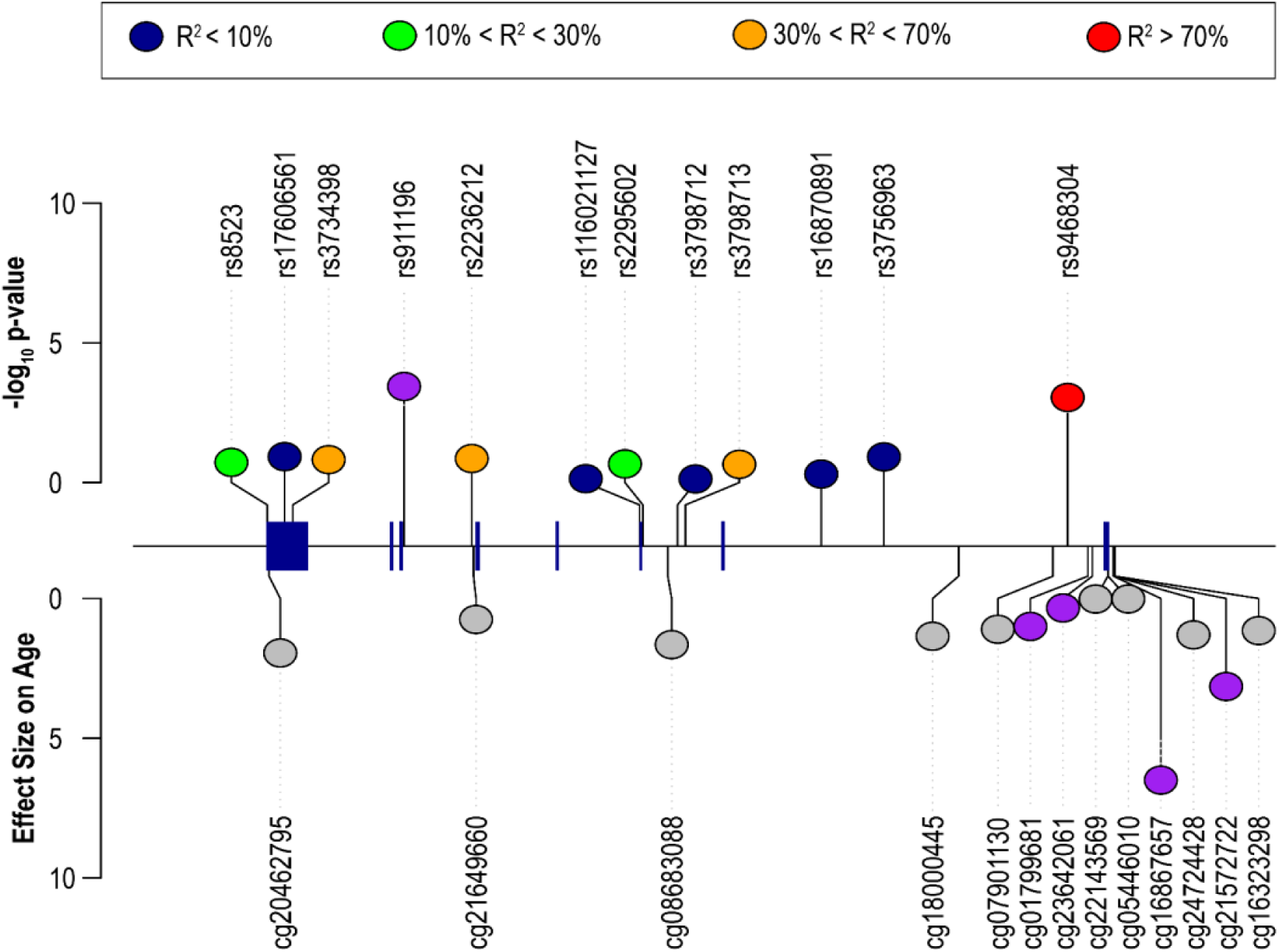
Lolliplot of variants correlated to age of onset of AMD in the ELOVL2 gene, relative to the gene structure. ***Top:*** The negative decadic P-Value of the correlation between the alleles and age of onset/diagnosis is depicted on the y-axis. The lead variant rs911196 is highlighted in purple and the correlated variants are color coded according to their genetic correlation (R^2^) to the lead variant. ***Bottom:*** CpGs found in the ELOVL2 locus and their effect on aging expressed as the slope (i.e., higher methylation in those sites was correlated to older age). In purple, CpGs significantly positively correlated with the G allele at rs911196 causing earlier onset of AMD are highlighted.

## DISCUSSION

In this work, we identify a potential therapeutic approach aimed at restoring health of aged retinas. First, we performed a comprehensive analysis of aging phenotypes and molecular pathways specific to the mouse aging retina. Concurrently, we systematically characterized both vision and molecular alterations in the retinas of animals deficient in ELOVL2 function. We found, on a molecular, metabolic, and functional level, that *Elovl2^C234W^* animals exhibit a significant acceleration of aging phenotypes. Drawing from our understanding of the metabolism of PUFAs in aging, we designed a treatment strategy involving the intravitreal injection of the elongation product of ELOVL2, 24:5n-3, and demonstrated its efficacy in enhancing vision in aged animals. We show that the proposed therapy reverses several aging phenotypes, including the accumulation of age-related sub-RPE deposits enriched in ApoE, C3, and C5b-9. Furthermore, at the molecular level, the treatment exhibited the capability to downregulate the innate immune response, complement activation, and oxidative stress response. Finally, our genetic analysis revealed genetic variants in the human *ELOVL2* locus that correlate with an earlier onset of intermediate AMD, indicating the translational relevance of the proposed 24:5n-3 treatment in patients afflicted with the disease.

The gene *ELOVL2* encodes a transmembrane enzyme involved in the synthesis of long (C22 and C24) n-3 and n-6 PUFAs (**Fig. 2A)**. Specifically, ELOVL2 elongates docosapentaenoic acid (DPA) (22:5n-3) to 24:5n-3, which is a precursor for very long chain PUFAs (VLC-PUFAs) and 22:6n-3, docosahexaenoic acid (DHA). Notably, total lipid extracts from the photoreceptors of AMD patients contain lower levels of DHA and VLC-PUFAs compared to those of age-matched normal donors^4^. The regulatory element of *ELOVL2* becomes increasingly methylated with age, a phenomenon observed across various tissues and animal species, including rodents and humans^15,17,39,40^. Notably, this reproducible direct relationship between the levels of methylation of the *ELOVL2* promoter and age has been shown to be one of the most reliable DNA methylation markers associated with chronological age of organisms^15,16,41–45^. To study the role of ELOVL2 in vision, we generated a homozygous mutant mouse, *Elovl2^C234W^*, that lacks the ability to convert the 22:5n-3 PUFA, DPA, to 24:5n-3^46,47^. In this study, our comprehensive lipidomic analysis unveiled striking similarities in the lipid composition of the retina and photoreceptor outer segments in aging and in response to *Elovl2* mutation. Our longitudinal studies on vision in aging and in *Elovl2^C234W^* animals show a significantly earlier onset of vision decline in the mutant retinas. Nevertheless, in both conditions, we did not detect any retinal degeneration or dysfunction in the overall visual cycle.

Aging is one of the major risk factors for developing AMD, and there is no cure or treatment to successfully slow down the disease progression in early or intermediate stages. Epidemiological studies have suggested that diets rich in n-3 PUFAs may be associated with lower occurrence of AMD, whereas low dietary intake of n-3 PUFAs may be associated with an opposite effect^48,49^. Two early surveys found that high plasma levels of n-3 PUFAs were correlated with decreased risk of AMD^50,51^. Nutritional supplementation with EPA (eicosapentaenoic acid) and DHA, however, has yielded inconclusive results. In two large prospective studies, the Age-Related Eye Disease Study-2 (AREDS2) and the nutritional AMD study (NAT-2), no significant difference was observed between oral supplementation with DHA and EPA *versus* placebo in the progression to wet AMD^12,52^. Interestingly, in the AREDS study, a large prospective study investigating factors influencing progression to advanced AMD, subjects with the highest self-reported intake of foods rich in n-3 LC-PUFAs were 30% less likely to develop central geographic atrophy (GA) and 50% less likely to develop AMD compared to those with the lowest self-reported intake^53^. Recent work from the Bernstein laboratory^10^ has suggested a potential explanation for this discrepancy, demonstrating that *Elovl4* mutant animals supplemented with oral 32:6n-3 VLC-PUFA, rather than DHA and EPA, exhibited a modest improvement in vision.

While several laboratories have undertaken studies with oral or dietary supplementation of PUFAs to prevent vision loss in aging or disease^10,11^, notable progress has only been achieved recently. In light of inconclusive results from EPA and DHA supplementation, focus has shifted to longer n-3 PUFAs. As mentioned above, a short-term feeding study reported the retinal incorporation of chemically synthesized 32:6n-3, a product of ELOVL4, and improvement of vision in wild-type and *Elovl4* mutant mice^10^. More recently, 8-week dietary supplementation of fish oil enriched in C24-C28 n-3 PUFAs in 9-month-old animals showed slightly improved ERG responses and visual performance, presenting promising alternatives to the DHA- and EPA-enriched diets^11^. Taken together, this data suggests that the beneficial impact of high fish intake observed in AREDS studies might be attributed to the requirement for PUFAs longer than EPA and DHA in the eye.

Here, following our analysis of physiology and lipid metabolism in both aged and *Elovl2^C234W^* animals, we decided to use a product of PUFA elongation facilitated by the ELOVL2 enzyme, namely 24:5n-3, in a direct supplementation strategy. We opted for the intravitreal route of delivery for this proof-of-concept study, anticipating that this will enable precise control over the levels of fatty acid delivered to the retina, as well as offer insights into potential therapeutic, in lieu of nutrition-based, strategies applicable in clinical settings. Our dose of lipid is significantly lower than any dose previously used in similar studies and was established as half of the absolute amount of 24:5n-3 in young (3-month-old) retinas. The single intravitreal injection did not induce adverse effects on vision, as tested in both young and old animals. In contrast, improvements in scotopic and photopic ERG recordings, VEP readings, and rod-mediated dark adaptation recovery underscored the beneficial effects of the treatment.

Lipidomic analysis following 24:5n-3 supplementation showed a low, but significant, increase in the VLC-PUFA-containing phospholipids in the retina. This outcome may be attributed to the low dose of the injected PUFA, and its intrinsic position in the PUFA elongation pathway as a substrate for DHA and VLC-PUFAs. We cannot, however, exclude the possibility that 24:5n-3 plays an additional role in the retina, as suggested by the study involving fish oil enriched in C24-C28 n-3 PUFAs^11^. Our transcriptome analysis revealed a robust downregulation of inflammation, including decreased activity of the complement pathway, in addition to downregulation of microglial phagocytosis and migration, hallmark processes affected in the aging retina^54–56^.

To further investigate the mechanisms underlying the improvement of vision in the aged animals following 24:5n-3 supplementation, we examined TFs that potentially regulate DEGs. Unsurprisingly, most of the detected TFs were downregulated and have been previously shown to be key regulators of retinal microglial sensome and pro-inflammatory genes in activated microglia^57^. Our data therefore underscores the significant role of microglia in retinal aging.

Specific look at the aging cerebellum pathway^33^ revealed significant downregulation of many cytokines, growth factors and other molecules, including ApoE, one of the main proteins accumulated in the aging eye and a risk factor for developing AMD. Our fluorescence immunostaining confirmed this striking downregulation, along with other factors such as C3, C5b-9 and HTRA1, all associated with aging and AMD risk. Taken together, our transcriptomic and immunostaining data suggest that the 24:5n-3 treatment induces a significant shift in tissue status towards a younger phenotype.

At the genetic level, several variants within the *ELOVL2* locus have been described. Specifically, variants within noncoding regions (introns and regulatory elements) have been associated with n-3 PUFA levels in plasma^58–60^ and colostrum^61^, as well as with risk of coronary artery disease^62^, Alzheimer’s disease^63^ and child cognition^61^. To date, despite numerous genome-wide studies, no *ELOVL2* mutations or variants have been detected that correlate with the risk of AMD. We hypothesized two possible explanations^24^: First, *ELOVL2* is an essential gene for population survival. It has been shown that heterozygosity of the gene in a mouse model causes infertility in C57BL/6 mice^47^. Therefore, variants that can be potentially correlated with the disease are rare and have yet to be discovered. Second, the downregulation of *ELOVL2* expression is not exclusive to AMD but is associated with aging. In this scenario, potential variants affecting the activity or expression of *ELOVL2* may correlate more with the onset of the disease rather than the disease itself. To address the latter possibility, we used genomic data from the International AMD Genomics Consortium and the UK Biobank and asked whether there are any variants in the *ELOVL2* genomic region that modify the onset of the disease. With this approach, several non-coding variants were identified within the gene region of ELOVL2. While the associated variants were not found to influence gene expression of ELOVL2 in retina or brain, the G allele of the lead variant rs911196 increases methylation of cg01799681, cg16867657, cg21572722 and cg23642061 in whole blood^64,65^. Those CpG sites are located near the ELOVL2 promotor and increased methylation at those sites is strongly associated with more advanced age^66^. Thus, the variant might influence ELOVL2 expression in certain cells (such as retinal cones, which would not be represented sufficiently in whole retina eQTL studies) and thus exert its effect on the onset of intermediate AMD. Interestingly, AREDS studies have indicated that higher intake of fish protects against progression of intermediate AMD, reinforcing the significance of our finding. To our knowledge, this data represents the first genetic evidence of the involvement of *ELOVL2* in AMD. We anticipate that future analyses will reveal genetic correlations between *ELOVL2* and other age-related diseases of the central nervous system.

We acknowledge several limitations of our study. First, this proof-of-concept work utilized intravitreal injection, which is an unlikely route of administration for preventing aging of the eye in patients. Future animal studies employing alternative administration routes, such as intraperitoneal injection, will be conducted to establish the effectiveness of the treatment. Additionally, the duration of treatment efficacy remains to be established. Furthermore, while we demonstrate that 24:5n-3 contributes to changes in levels of VLC-PCs and other phospholipids, we take into consideration that this fatty acid may exert protective effects through additional mechanisms. Investigations utilizing labeled 24:5n-3 should address these questions.

In sum, our work demonstrates that targeted delivery of a product of ELOVL2 to the eye can reverse age-related molecular and functional changes. We have discovered a genetic connection between the *ELOVL2* locus and the onset of intermediate AMD. Our work is paving the way towards developing novel lipid-based therapeutic strategies aimed at preserving visual health in aging populations.

## ACKNOWLEDGEMENTS

The authors would like to thank Vincent Du Vinh-Mai, Mykala Garner, and Zach Pope for invaluable help in this project. This work was supported by Edward N. and Della L. Thome Memorial Foundation, BrightFocus foundation and NIH grants U01EY034594 (DSK) and R01EY035137 (VJK). The authors acknowledge support to the Gavin Herbert Eye Institute at the University of California, Irvine from an unrestricted grant from Research to Prevent Blindness and from NIH core grant P30 EY034070. ET is supported by the Visual Sciences Training Program (T32EY032448) and F30EY035146. ATF is supported by the National Science Center, Poland (2019/34/E/NZ5/00434).

## ONFLICT OF INTEREST

DS-K is a scientific advisor of Visgenx, Inc. The remaining authors declare no conflict of interest.

## MATERIALS AND METHODS

### Electroretinography (ERG)

Animals were dark-adapted overnight and anesthetized by IP injection of ketamine (100 mg/kg) and xylazine (4 mg/kg). During recordings, a heating pad-maintained body temperature at 37–38°C. The eyes were dilated using 1% atropine sulfate (rod dark adaptation experiments) or 1% tropicamide and 2.5% phenylephrine (regular scotopic and photopic recordings). The eyes were lubricated with a corneal gel. The reference electrode needle was inserted under the skin at the skull. The responses for eleven stimuli of increasing intensities were recorded and averaged. The a- and b-wave responses were plotted from the averaged ERG waveform. Scotopic and photopic ERG recordings were performed using commercially available platform (Celeris, Diagnosys LLC). For photopic recordings mice were kept in a lighted vivarium for 10 minutes prior to experiment. M-cone and S-cone specific function was measured. Stimulation was performed by the means of alternating UV and green light with flashes of increasing intensities. Green light stimulation had steps of increasing intensities of −0.5, 0.5, 1.5, and 2.5 log cd·s/m2. UV light stimulation had increasing intensities of −1, 0, 1, and 2 log cd·s/m2. The responses for treated and non-treated eyes were grouped respectively and averaged together. The a- and b-wave tracings represent the averaged ERG waveform. The data was analyzed with Espion v.6 software (Diagnosys). GraphPad Prism was used for graph preparation and statistical analysis.

### Dark Adaptation measurements

For rod dark adaptation experiments, full-field ERGs were recorded using a UTAS BigShot apparatus (LKC Technologies). Scotopic responses to calibrated green (530 nm) LED light were recorded from both eyes. Measurements from several trials were averaged and the intervals between trials were adjusted so that responses did not decrease in amplitude over the series of trials for each step.

Rod ERG a-wave fractional flash sensitivity (*S_f_*) was calculated from the linear part of the intensity-response curve, as follows:

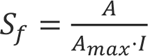

where *A* is the amplitude of the rod a-wave dim flash response, *A_max_* is the maximum amplitude of the rod a-wave response for that eye (determined at 23.5 cd·s m^-2^), and *I* is the flash strength. The maximal response and sensitivity of rods were first determined in the dark. To monitor the post-bleach recovery of *A_max_*and *S_f_*, more than 90% of rhodopsin was bleached with a 35-sec exposure to 520-nm LED light focused at the surface of the cornea. The fraction of a bleached pigment was calculated with the following equation:

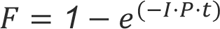

where *F* is the fraction of rhodopsin bleached, *t* is the duration of the light exposure (s), *I* is the bleaching light intensity of 520-nm LED light (1.3 x 10^8^ photons µm^-2^ s^-1^), and *P* is the photosensitivity of mouse rods at the wavelength of peak absorbance (5.7 x 10^-9^ µm^2^)^67^. Mice were re-anesthetized once after 30 min post-bleach with a lower dose of ketamine (∼1/3 of the initial dose). If needed, a small drop of PBS solution was gently applied to their eyes with a plastic syringe to protect them from drying and to maintain contact with the recording electrodes. For each time point, the *A*_max_ and *S_f_* were normalized to the corresponding *A*_max_^DA^ and *S_f_*^DA^ values.

### Optomotor responses (OMR)

Optomotor responses (OMRs) were recorded using a commercially available OMR setup. (PhenoSys GmbH, Berlin, Germany). The software automatically tracks animal head movement in direction of rotating gratings stimulus (optomotor reflex) and calculates correct/incorrect tracking behavior presented as OMR (optomotor) index. The mouse was placed on an elevated platform in OMR arena. Gratings stimuli (rotating at 12°/s) were presented for ∼12 min per trial. The spatial frequency of the gratings was set at 0.2 cycles per degree of visual angle. The stimulus was presented at differing contrasts between the light and dark stripes: 3, 6, 10, 12.5, 15, 17.5, 20, 25, 37.5, 50, 75, 100 contrast. The stimulation at each contrast level lasted for 60 s. For the scotopic part of the experiment (nighttime light levels) the animals were dark-adapted for 12h before the experiments. The OMR arena was dimmed using 4 neutral density filters in front of the stimulus displays (to ∼0.03 lux). Results for 100% contrast were excluded due to adjustment (mice adjustment to the system). OMR results that yielded correct/incorrect ratio lower than 0.8 were excluded. For each mouse, the results from several trials were averaged for analysis. Each data point represents the average of all mice tested. Graphpad prism was used for statistical analysis and to generate curves.

### Neurophysiology (Visual Evoked Potential - VEP)

Animals were initially anesthetized with 2% isoflurane in a mixture of N_2_O/O_2_ (70%/30%) then placed into a stereotaxic apparatus, followed by analgesia (Flunixin Meglumine, 2.5 mg/kg SC every 8 hours) and local subcutaneous injection of lidocaine (0.5%). A small, custom-made plastic chamber was secured to the exposed skull using dental acrylic. After one day of recovery, re-anesthetized animals were placed in a custom-made hammock, maintained under isoflurane anesthesia (1-2% in N_2_O/O_2_), a craniotomy was performed, and multiple single tungsten electrodes were inserted into V1 layers II-VI. Following electrode placement, the chamber was filled with sterile agar and sealed with sterile bone wax. Animals were then sedated with chlorprothixene hydrochloride (1 mg/kg; IM) and kept under light isoflurane anesthesia (0.2 – 0.4% in 30% O_2_) throughout the recording procedure. EEG and EKG were monitored throughout, and body temperature was maintained with a heating pad (Harvard Apparatus, Holliston, MA).

Data was acquired using a multi-channel Scout recording system (Ripple, UT, USA). Local field potentials (LFP) from multiple locations at matching cortical depths were band-pass filtered from 0.1 Hz to 250 Hz and stored along with spiking data at a 1 kHz sampling rate. LFP signal was aligned to stimulus time stamps and averaged across trials for each recording depth in order to calculate visually evoked potentials (VEP)^68^. Single neuron spike signals were band-pass filtered from 500 Hz to 7 kHz and stored at a 30 kHz sampling frequency. Spikes were sorted online in Trellis (Ripple, UT, USA) while performing visual stimulation. Visual stimuli were generated in Matlab (Mathworks, USA) using Psychophysics Toolbox and displayed on a gamma-corrected LCD monitor (55 inches, 60 Hz; 1920 x 1080 pixels; 52 cd/m^2^ mean luminance). Stimulus onset times were corrected for monitor delay using an in-house designed photodiode system^69^. Visual responses were assessed according to previously published methods^69,70^. For recordings of visually evoked responses, animals were tested with 100 repetitions of a 500 ms bright flash of light (105 cd/m^2^).

### Optical coherence tomography (OCT)

Mice were anesthetized via intraperitoneal injection with ketamine/xylazine (10mg/kg), followed by pupil dilation using 1% tropicamide and 2.5% phenylephrine. Optical coherence tomography (OCT) was performed using a Bioptigen spectral-domain OCT device (Leica Microsystems Inc., Buffalo Grove, IL), capturing four frames of OCT b-scan images from a series of 1200 a-scans. Outer nuclear layer (ONL) thickness was assessed 500 µm from the optic nerve head (ONH) at the nasal, temporal, superior, and inferior quadrants of each eye, and these values were averaged to determine the average ONL thickness.

### POS isolation

Photoreceptor outer segments (POS) were isolated by OptiPrep™ density gradient centrifugation described previously with minor modification^71^. Briefly, 1 retina was placed in Ringer’s solution (10 mM HEPES (pH 7.4), 130 mM NaCl, 3.6 mM KCl, 12 mM MgCl2, 1.2 mM CaCl2, and 0.02 mM EDTA] containing 8% OptiPrep™ and vortexed for 1 min.The samples were centrifuged at 200 x g for 1 min at 4 °C and the supernatant containing the ROS was collected gently. The vortexing and sedimentation sequence was repeated six times. The supernatant was combined for lipid extraction.

### Intravitreal Injections

All animals were anesthetized with Isoflurane; their eyes were anesthetized with proparacaine (0.5%, Bausch-Lomb) and followed by dilation of pupils with a drop of tropicamide (1%, Alcon Laboratories) and of phenylephrine (2.5%, Akorn Pharmaceuticals, Lake Forest, IL). All drops were applied with pipette. Injections were performed under a surgical microscope (Zeiss). First, a drop of Gonak (Akorn Pharmaceuticals, Lake Forest, IL) was applied to the corneal surface for better visibility and prevention of back-leak from the eye. Next, a 33-gauge hypodermic needle was used to create a tunnel reaching into vitreous. A 33-gauge blunt-end needle (World Precision Instruments), connected to an RPE-KIT (World Precision Instruments) by SilFlex tubing (World Precision Instruments; SILFLEX-2), was advanced through the tunnel. Each mouse received 1 μl injection of a compound per eye.

### Lipidomic analysis

#### Lipid extraction

Lipid extractions were performed according to the methodology of Bligh and Dyer^18^. In brief, the tissue was homogenized in 200 μL water, transferred to a glass vial, and 750 μL 1:2 (v/v) CHCl_3_: MeOH was added and vortex well. Then 250 μL CHCl_3_ was added and vortex well. Finally, 250 μL ddH_2_O was added and vortex well. The samples were centrifuged at 3000 RPM for 5 min at 4 °C. The lower phase was transferred to a new glass vial and dried under nitrogen stored at -20 °C until subsequent lipid analysis.

#### LC-MS/MS

Separation of lipids was performed on an Accucore C30 column (2.6 μm, 2.1 mm × 150 mm, Thermo Scientific). The Q Exactive MS was operated in a full MS scan mode (resolution 70,000 at m/z 200) followed by ddMS2 (17,500 resolution) in both positive and negative mode. The AGC target value was set at 1E6 and 1E5 for the MS and MS/MS scans, respectively. The maximum injection time was 200 ms for MS and 50 ms for MS/MS. HCD was performed with a stepped collision energy of 30 ± 10% for negative and 25%, 30% for positive ion mode with an isolation window of 1.5 Da.

#### Data analysis and post-processing

Data were analyzed with LipidSearch 4.2.21 software. Only peaks with molecular identification grade: A or B were accepted (A: lipid class and fatty acid completely identified or B: lipid class and some fatty acid identified). The relative abundance of each lipid species was obtained by normalization to the total lipids intensity. VLC-PUFAs incorporated into PCs were analyzed using Lipid Data Analyzer with a customized database. Significantly changed lipid species (FC>1.5. P-value<0.05) were submitted to Lipid Ontology (LION) for lipid ontology analysis. Data visualization was performed on Prism 7 software (GraphPad Software, Inc.).

#### FA analysis

Separation of VLC-PUFAs was achieved on an Acquity UPLC® BEH C18 column (1.7 μm, 2.1 mm × 100 mm, Waters Corporation). The Q Exactive MS was operated in a full MS scan mode (resolution 70,000 at m/z 200) in negative mode. For the compounds of interest, a scan range of m/z 250–800 was chosen. The identification of fatty acids was based on retention time and formula.

### RNA sequencing

#### Sample collection and preparation

Fresh retina tissue was dissected from each mouse eye. Total RNA was extracted from the retinas using the RNeasy Plus Mini Kit (Qiagen) following the manufacturer’s instructions. RNA quantity and quality were assessed using the Qubit RNA HS Assay kit (Thermo Fisher Scientific) and the Agilent Bioanalyzer RNA 6000 Pico kit (Agilent Technologies), respectively.

#### Library construction and sequencing

RNA sequencing libraries were prepared using the Illumina TruSeq RNA Library Prep kit. Paired-end sequencing was performed on the NovaSeq 6000 System using the Flow Cell Type S4, generating paired-end reads with a length of 100 base pairs, with approximately 30 million reads per sample.

#### Bioinformatics analysis

Raw sequencing reads were preprocessed to remove adapter sequences and low-quality bases using Trimmomatic. Cleaned reads were mapped to the mm10 assembly mouse reference genome using the aligner tool HISAT2. Gene-level expression counts were quantified using featureCounts. Differential expression analysis was performed using DESeq2, and genes with an adjusted p-value < 0.05 and |fold change| > 1.5 were considered differentially expressed.

### Single-nuclei RNA sequencing (snRNA-seq)

#### Sample collection and preparation

Retinas (n=3 per age group) were dissected and snap-frozen in liquid nitrogen. Deep frozen samples were sent to Active Motif for further processing. Active Motif is a 10x Genomics Certified Service Provider of Single-Cell Multiome to measure genome-wide gene expression & open chromatin. Samples were processed according to the Illumina “Isolation of Nuclei for Single Cell RNA Sequencing & Tissues for Single Cell RNA Sequencing” procedure and subjected to the snRNAseq following the Illumina protocol.

#### Quality control, integration and clustering

91bp sequencing reads were generated by Illumina sequencing. Reads were mapped to the (reference genome) and counts at features were identified using the Cell Ranger for Single Cell Gene Expression software with default parameters (mkfastq and count functions). Quality control was performed, and cellular barcodes that matched the following three criteria were kept: number of unique molecular identifiers (UMIs) within three median absolute deviations (MADs) of the population median, number of expressed genes within three MADs of the population median, and the percentage of reads mapping to mitochondrial genes under 15%. We detected the expression of approximately 1000 genes per nucleus in each sample. Downstream analysis was performed using the Seurat (v4.3.0) R package. Data were normalized using Seurat’s LogNormalize function and optimal cluster resolution was determined using the clustering-tree method^72^.

To visualize cells onto a two-dimensional space, we performed Uniform Manifold Approximation and Projection (UMAP). Cell identities were assigned using known markers established in previous studies and markers identified in our study. The Wilcoxon Rank Sum test was used to perform differential gene expression analysis between the clusters. Genes with a p-value of less than 0.05 and a log fold change greater than 1 were considered as differentially expressed. Additionally, the Wilcoxon Rank Sum test was used to perform differential gene expression analysis between young and aged conditions within each cell type. Genes with a p-value of less than 0.05 and a log fold change greater than 0.1 were considered as differentially expressed. The FeaturePlot of Elovl2 expression levels was normalized to TBP.

### Cell-cell communication analysis

Cell-cell communication analysis was performed using the CellChat package (v.2.0.0). Only cells in cones, rods and Müller glia were used to infer the cell–cell communication. We first applied CellChat to young (3-month-old) and aged (18-month-old) datasets separately to infer cell-cell communication followed by comparison analysis via merging the CellChat objects from young and aged datasets. In brief, we followed the official workflow and loaded the normalized counts into CellChat and applied the preprocessing functions ‘identifyOverExpressedGenes’, ‘identifyOverExpressedInteractions’ and ‘projectData’ with standard parameters set. For the main analyses the core functions ‘computeCommunProb’, ‘computeCommunProbPathway’ and ‘aggregateNet’ were applied using standard parameters.

### Gene set enrichment analysis (GSEA)

Gene Set Enrichment Analysis (GSEA) was conducted using the GSEA software (v4.2.3)^73^. Differentially expressed genes identified by DESeq2 were ranked using the signal-to-noise ratio rank metric. Enrichment analysis was performed using predefined gene sets from databases such as Gene Ontology (GO), Molecular Signatures Database hallmark gene set, Reactome, and Wikipathways^74–77^. Statistical significance of enrichment was determined using a weighted (*p*=1) scoring calculation scheme with 1000 permutations. Gene sets with size larger than 500 and smaller than 5 were excluded from the analysis.

### Metascape gene enrichment and functional analysis

Differentially expressed genes (DEGs) (adjusted p-value < 0.05, |fold change| > 1.5) were inputted into the Metascape platform, where pathway and process enrichment analysis were carried out with selected ontology sources including GO Biological Processes, KEGG Pathway, Reactome Gene Sets, and WikiPathways. Terms with a p-value < 0.01, a minimum count of 3, and an enrichment factor > 1.5 (the enrichment factor is the ratio between the observed counts and the counts expected by chance) were collected and grouped into clusters based on their membership similarities. Then, to further capture the relationships between the terms, a network plot was constructed for a subset of enriched terms, where terms with a similarity > 0.3 were connected by edges. This network was visualized using Cytoscape (v3.10.0), where each node represents an enriched term and is proportional to the number of input genes in each term and is colored by its Cluster ID or p-value.

### Transcription factor enrichment analysis

Transcription factor (TF) enrichment analysis was performed using the ChEA3 (ChIP-X Enrichment Analysis 3) web tool for differentially expressed genes (DEGs) (adjusted p-value < 0.05, |fold change| > 1.5). TFs were ranked using the Mean Rank metric, which averaged integrated ranks across gene set libraries. A global transcription co-expression network was constructed from GTEx expression data for the top 20 TFs and clustered by GO enrichment term. Protein-protein interaction analysis was performed for the top 20 associated TFs using the Search Tool for the Retrieval of Interaction Genes/Protein (STRING) database. After filtering out TFs with low expression levels (normalized read count DESeq2 baseMean < 10), protein-protein interactions were visualized in Cytoscape (v3.10.0) as nodes and edges, and each node was colored by the difference in mean Z-score.

### Bisulfite sequencing for DNA methylation analysis

The methylation profile of the DNA was determined using PCR amplification followed by DNA Sanger sequencing. Methylated DNA was converted using EZ DNA Methylation-Direct™ Kit (Zymo Research, USA) following the instructions of the manufacturer. The converted DNA was amplified using primers that do not overlap with CpGs. The amplified product was sequenced to detect the methylation status of individual cytosines within the amplified region. The methylation percentage at each CpG site was computed as the ratio of methylated cytosines to the total level signal of cytosines.

### Retinoid analysis

Animals were dark-adapted overnight, and eyes were enucleated following euthanasia under dim red light and flash-frozen in liquid nitrogen. Retinoids were extracted, derivatized, and separated on an Agilent Rx-SIL HPLC column (Neta Scientific) using a two-step normal-phase HPLC protocol run on an Agilent 1260 Infinity II series LC instrument as described previously^22,78^. Less polar retinyl esters are first separated and eluted in a mixture of 0.6% ethyl acetate:99.4% hexanes at a flow rate of 1.4 mL/min, and more polar retinal-oximes and retinols are subsequently separated and eluted in a mixture of 10% ethyl acetate:90% hexanes at a flow rate of 1.4 mL/min.

### Retina cryosection and immunohistochemistry

Mouse eyes were enucleated following euthanasia and fixed in 4% paraformaldehyde (PFA) in phosphate-buffered saline (PBS, pH 7.4) for 2 hours at 4°C. The cornea, lens, and vitreous were removed, and the eyecups were fixed in 4% PFA overnight at 4°C. Following fixation, eyecups were cryoprotected by immersion in a sucrose gradient (10% and 20% sucrose for 1 hour at room temperature, and 30% sucrose overnight at 4°C). Eyecups were embedded in Tissue-Tek OCT (Sakura, Torrance, CA) and frozen on a conductive metal block placed on dry ice. Cryosectioning was performed using a cryostat, and retina sections were cut into 10 µm-thick serial sections.

For immunohistochemistry, sections were blocked in 0.3% TritonX-100 in 5% BSA for 1 hour at room temperature to minimize nonspecific binding. Sections were then incubated with primary antibodies diluted in 0.1% TritonX-100 in 5% BSA (see Table 1) overnight at 4°C. Following three washes with PBS, sections were incubated with fluorescently-labeled secondary antibodies diluted in 0.1% TritonX-100 in 5% BSA (see Table 1) for 1 hour at room temperature. Following three washes with PBS, nuclei were counterstained with Hoescht 33342 (Thermo), and sections were mounted using ProLong Gold Antifade (Thermo). Immunostained sections were imaged on a Keyence All-in-One Fluorescence microscope (BZ-X810, Keyence, Itasca, IL) at 40X and 100X magnification.

**Table 1.** Catalog numbers of antibodies and RNA probes used in experiments.

| Antibody/Probe Name | Species | Dilution | Company (Catalog #) |
| --- | --- | --- | --- |
| C3 | rat | 1:50 | Santa Cruz (sc-58926) |
| APOE | mouse | 1:50 | Santa Cruz (sc-13521) |
| C5b-9 | mouse | 1:50 | Santa Cruz (sc-66190) |
| PKCa | mouse | 1:200 | Santa Cruz (sc-8393) |
| GS | rabbit | 1:200 | Sigma (HPA007316) |
| HTRA1 | rabbit | 1:50 | Santa Cruz (sc-50335) |
| Elovl2 | mouse | - | Advanced Cell Diagnostics (542711-C3) |
| Elovl4 | mouse | - | Advanced Cell Diagnostics (835111-C1) |
| Elovl5 | mouse | - | Advanced Cell Diagnostics (868931-C2) |

### RNAscope

In situ hybridization was performed using the RNAscope® Multiplex Fluorescent Assay v2 (ACD Diagnostics). Probes used were designed by the manufacturer (see Table 1). Briefly, fresh frozen histologic sections of mouse eyes were pretreated per manual using hydrogen peroxide and target retrieval reagents including protease IV. Probes were then hybridized according to the protocol and then detected with TSA Plus® Fluorophores fluorescein, cyanine 3, and cyanine 5. Sections were mounted with ProLong Gold Antifade (Thermo Fisher) and imaged (Keyence BZ-X810).

### Proteomic analysis

Mouse retinas were harvested and suspended in a Urea buffer containing 8 M Urea, 0.1 M Tris-HCl (pH 8.5), protease inhibitor cocktail (cOmplete™ Protease Inhibitor Cocktail, Roche). The suspension was sonicated on ice for 4 min, followed by centrifugation at 12,000 g for 10 min at 4 °C. The supernatant was collected and digested by the filter-aided sample preparation (FASP) method^79^. Briefly, the supernatant was transferred into a spin filter column (30 kDa cutoff). Proteins were reduced with 10 mM DTT for 1 hr at 56 °C, and alkylated with 20 mM iodoacetic acid for 30 min at room temperature in the dark. Next, the buffer was exchanged with 50 mM NH_4_HCO_3_ by washing the membrane three times.

For quantification of Rho, PED6A, PDE6B, GNAT1, GBB1, stable isotope–labeled (SIL) peptides (listed in **Table 1**) were spiked into protein samples and then free trypsin was added into the protein solution at a trypsin to protein ratio of 1:50 and incubated overnight at 37 °C. The tryptic digests were recovered in the supernatant after centrifugation, and an additional wash with water. The combined supernatants were vacuum-dried and then dissolved in 20 μL 0.1% FA in H_2_O.

#### Mass spectrometry data acquisition

Proteomics data were acquired via LC-MS/MS using a Vanquish (Thermo Fisher Scientific), coupled in-line with a Q Exactive mass spectrometer (Thermo Fisher Scientific) with an ESI source. Mobile phase A was composed of 0.1% formic acid (FA) in water, and mobile phase B was composed of 0.1% formic acid (FA) in acetonitrile. The total flow rate was 0.4 mL min^-1^.

For parallel analysis monitoring (PRM), peptides were separated over a 25 min gradient on an Acquity UPLC® BEH C18 column (1.7 μm, 2.1 mm × 100 mm, Waters Corporation). MS2 scan parameters were set to select the m/z ratio of endogenous and the corresponding SIL peptides of PED6A, PDE6B, GNAT1, GBB1, GBB2 with defined elution time windows (Table S2). MS1 scans were acquired at the m/z range of 300–1000, mass resolution of 70,000, automatic gain control (AGC) target of 1e6, and maximum ion injection time of 50 ms. The PRM scans were acquired at a resolution of 17,500, AGC target value of 1e5, maximum ion injection time of 50 ms, isolation window of 2.0 m/z, and the HCD energy was 25%.

#### Data Processing

For PRM quantification, Skyline was used for processing the data. Peak picking and peak boundaries were carried out by Skyline and manually adjusted based on the overlapping precursor and product peaks.

**Table 2.** Sequences of selected SIL peptides, purity, concentration of stock solution and spiking Solution.

| Proteins | Sequences of SIL peptides | Purity/% |
| --- | --- | --- |
| PDE6A | F{Leu(13C6,15N)}DSNIGFAK | 86.8 |
| PDE6B | FLLAEESLN{Ile(13C6,15N)}YQNLNR | 86.9 |
| GNAT1 | EMSDIIQ{Arg(13C6,15N4)} | 82.0 |
| GBB1 | LLLAGYDDFN{Cys(CAM)}N{Val(13C5,15N)}WDALK | 77.8 |
| GBB2 | TFVSGA{Cys(CAM)}DAS{Ile(13C6,15N)}K | 84.8 |
| Rho | NP{Leu(13C6,15N)}GDDDASATASK | 83.9 |

### Analysis of *ELOVL2* variants and AMD

To study the role of *ELOVL2* genetics in AMD risk, we computed the correlation between those variants and age at disease onset or diagnosis. First, we identified incident AMD cases from the UK Biobank (the data was accessed and analyzed under the UK Biobank project ID 73446). Detailed ophthalmological grading is not available for the majority of the UK Biobank participants. Therefore, we leveraged the general practice read codes, as previously described^80^ to ascertain incident AMD patients. In total, we found 1,309 unrelated European individuals that developed any type of AMD in the UK Biobank after recruitment. In this case-only analysis, we used linear regression with age at onset as the outcome and each variant separately as exposure. The analyses were adjusted by sex, BMI, smoking status and the first ten principal components of ancestry. To replicate our findings, we extracted all non-imputed variants from the Illumina HumanCoreExome in the genomic region of *ELOVL2* (chr6: 10980992 – 11044624) for unrelated and European participants with intermediate AMD from a recent GWAS (n=2,407) conducted by the International AMD Genomics Consortium^38^. Similar to the UK Biobank analyses, we used linear regression adjusted for study, sex and the first five principal components of ancestry. The resulting slopes and standard errors from both cohorts were then meta-analyzed with the REML algorithm implemented in the *rma* function from the *metafor* package^81^ in R. The association results were plotted as a lolliplot, implemented in the *trackViewer* package in R^82^.

## Supplementary

**Fig. S1.**
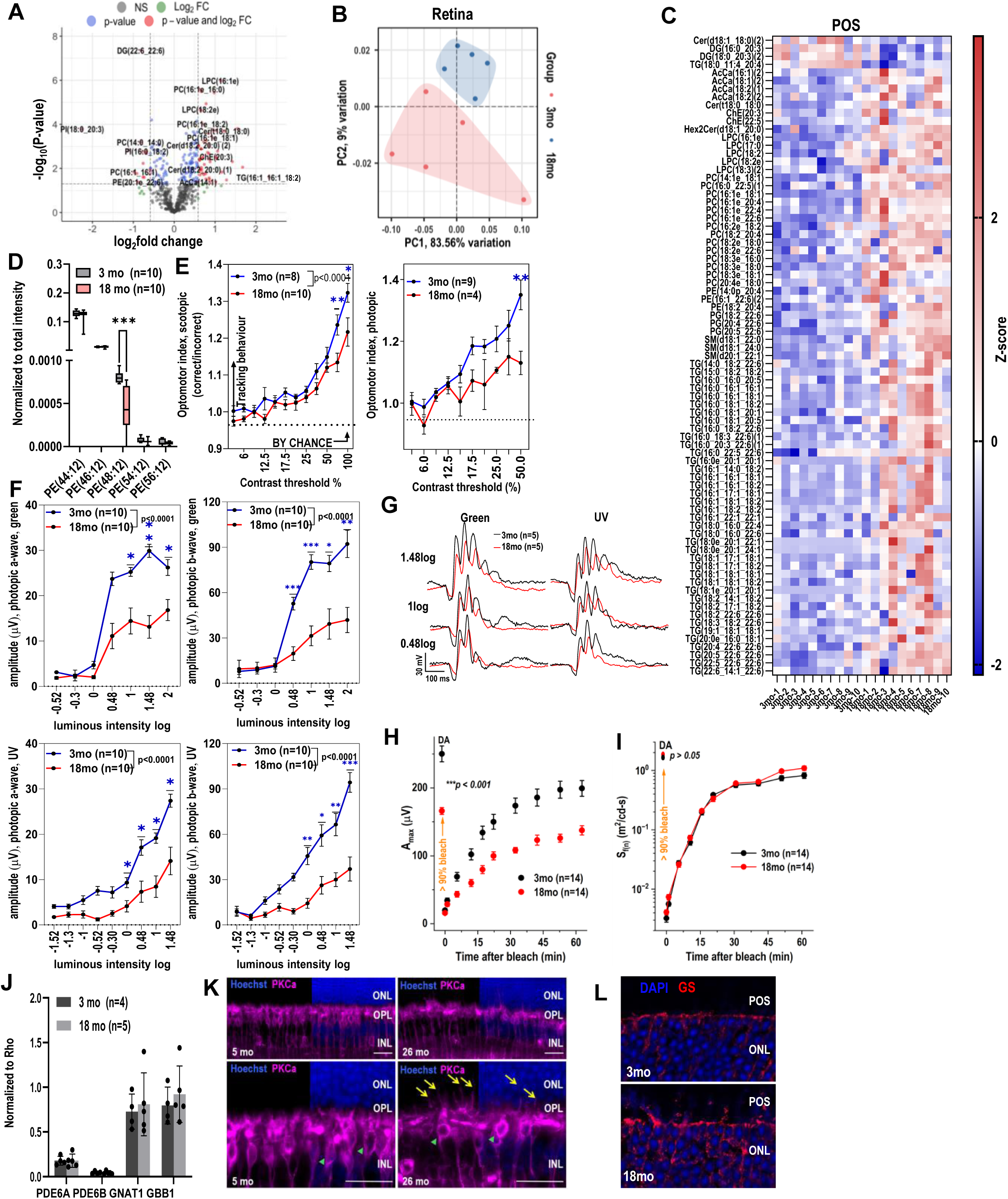
Age-related decreased VLC-PUFA levels in the retina is associated with vision decline. (**A**) Untargeted lipidomics analysis of young (3-month-old) and aged (18-month-old) retinas (n=5) showed 15 significantly down regulated lipids and 53 up-regulated lipids in aged retina (FC > |1.5|, pval<0.05). **(B)** Significantly changed lipids (FC>|1.5|, p<0.05) exhibited distinct clustering between young (3-month-old) and aged (18-month-old) retinas by principal component analysis (PCA). (**C**) Significantly changed lipid species in aging POS. (**D**) Among the analyzed VLC-PEs, PE (48:12) was significantly downregulated in aging POS (n=10, *** = p<0.001). (**E**) Decreased contrast sensitivity and (**F**) photopic electroretinogram (ERG) responses in aged (18-month-old) animals compared to young (3-month-old) animals (* = p<0.05, ** = p<0.01, *** = p<0.001). (**G**) Photopic ERG waveforms in response to UV and green flash in 3- and 18-month-old mice. Three light stimulus intensities are presented: 0.48log[cd s m-2], 1log[cd s m-2] and, 1.48log [cd s m-2]. Each waveform is an average of n=5 animals in representative groups. (**H**) Decreased recovery of the averaged A_max_ in aged mice (17-month-old) compared to young mice (3.5-month-old mice). (**I**) The recovery of rod-driven ERG a-wave sensitivity (S_f_) following the same bleach was not compromised in older animals. (**J**) Relative amounts of proteins involved in phototransduction normalized to Rho remained unchanged in aging retina. (**K**) Immunofluorescence staining with rod bipolar-specific antibody (PKCa) showed age-related morphological changes in outer plexiform layer including overgrowth of dendrites into the outer nuclear layer. (**L**) Immunofluorescence staining with glutamine synthetase antibody (GS) showed age-related Muller cell processes passing outer limiting membrane.

**Fig. S2.**
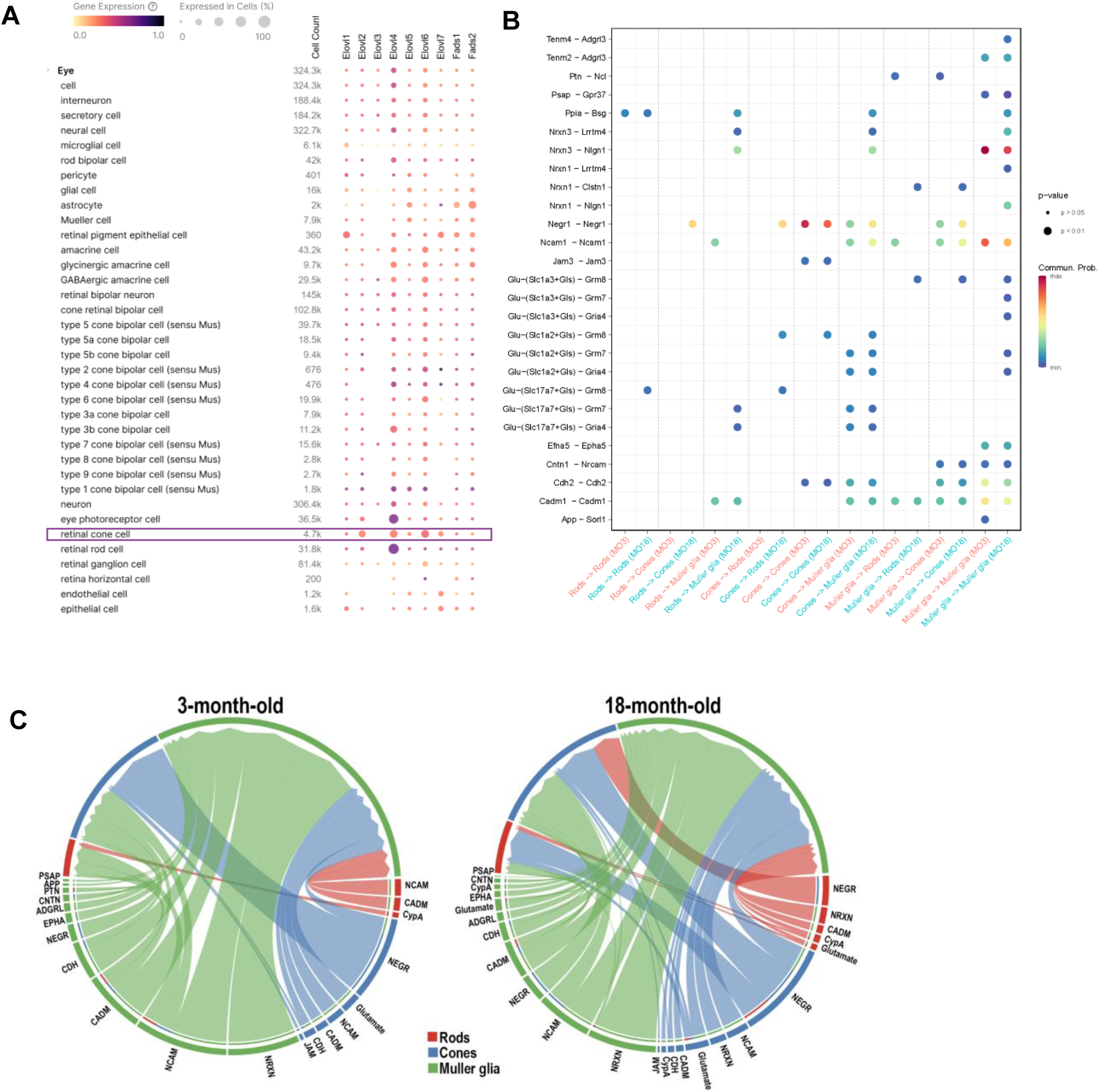
scRNA-seq analysis of genes involved in VLC-PUFAs biosynthesis pathway. (**A**) CZ CELLxGENE Discover visualization of expression of genes involved in VLC-PUFAs biosynthesis pathway in different cell types of the eye (**B**) Visualization of the communication probabilities mediated by ligand-receptor (L-R) pairs from specific cell groups to other cell groups in the young (3-month-old) and old (18-month-old) retinas. The x-axis represents the source-target cell group pairs, while the y-axis lists the different L-R pairs. The color intensity of the dots corresponds to the communication probability. The size of the dots represents the significance of the interactions CellChat interaction (**C**) Visualization of the signaling pathways mediating interactions in young and old retinas. Each segment along the circle represents a cell group, and the connecting arcs indicate the interactions between these groups. The color of the arcs corresponds to the source cell group. The thickness of the arcs indicates the interaction strength of the signaling pathway.

**Fig. S3.**
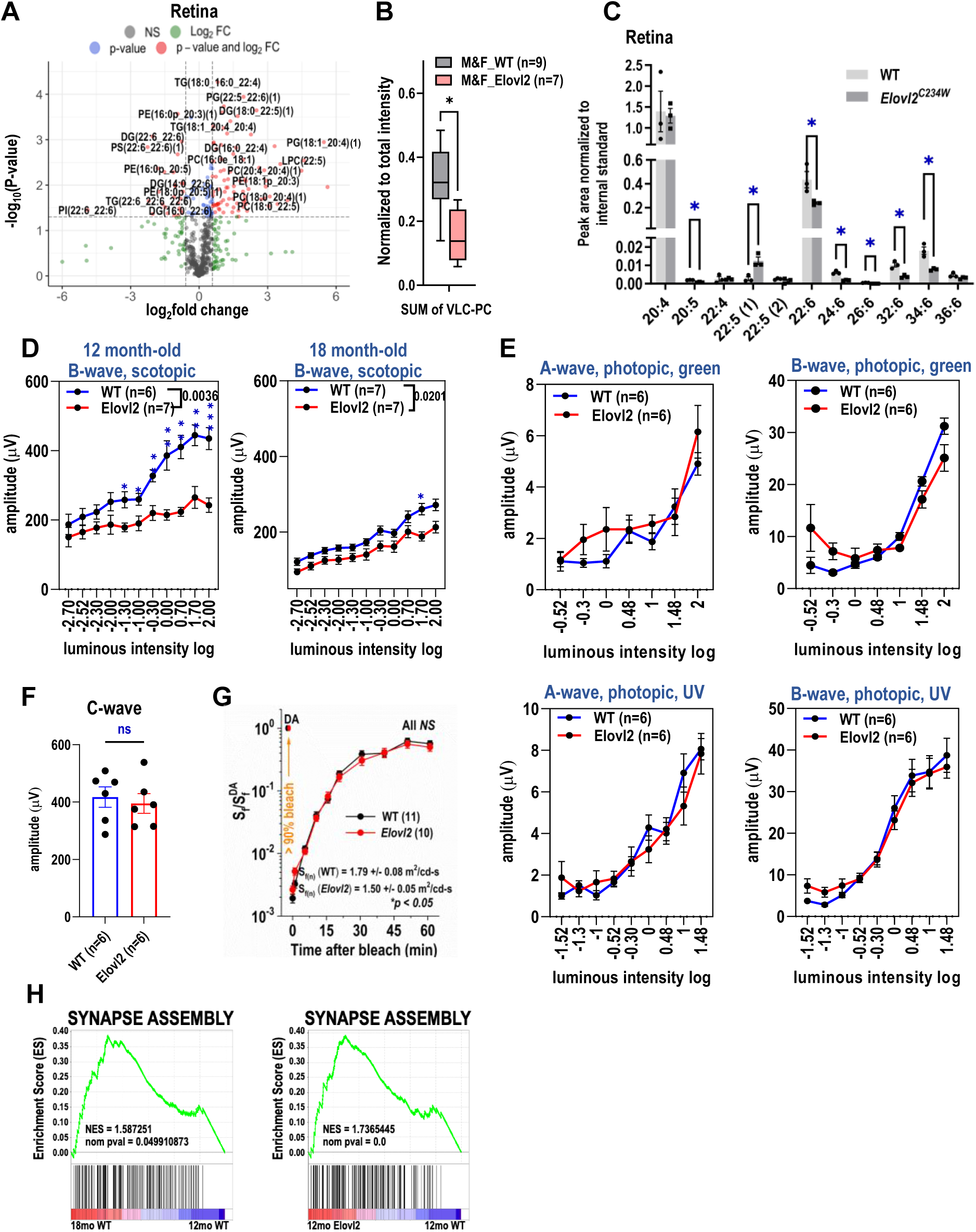
Disturbed lipid composition in Elovl2^C234W^ mouse retinas is correlated with vision loss. (**A**) Untargeted lipidomics analysis of 18-month-old wildtype and Elovl2^C234W^ retinas revealed 112 significantly changed lipids (FC>1.5, p-value <0.05). (**B**) Decreased total levels of VLC-PC in Elovl2^C234W^ retinas compared to age-matched wildtype retinas (* = p<0.05). (**C**) Free fatty acids analysis showed decreased DHA and VLC-PUFAs in Elovl2^C234W^ retinas (n=3, * = p<0.05). (**D**) Reduced scotopic b-wave electroretinogram (ERG) responses in 12- and 18-month-old Elovl2^C234W^ mice (* = p<0.05, ** = p<0.01, *** = p<0.001). (**E**) Photopic a- and b-wave, **(F)** scotopic c-wave ERG responses and (**G**) the recovery of rod-driven S_f_ remained unchanged in 12-month-old *Elovl2^C234W^* mice compared to age-matched wildtype mice (n=6). **(H)** GSEA enrichment plots for synapse assembly (GO: 0007416) pathway showed similar enrichment in 12-month-old Elovl2^C234W^ vs. age-matched wildtype retinas and 18-month-old wildtype vs. 12-month-old wildtype retinas.

**Fig. S4.**
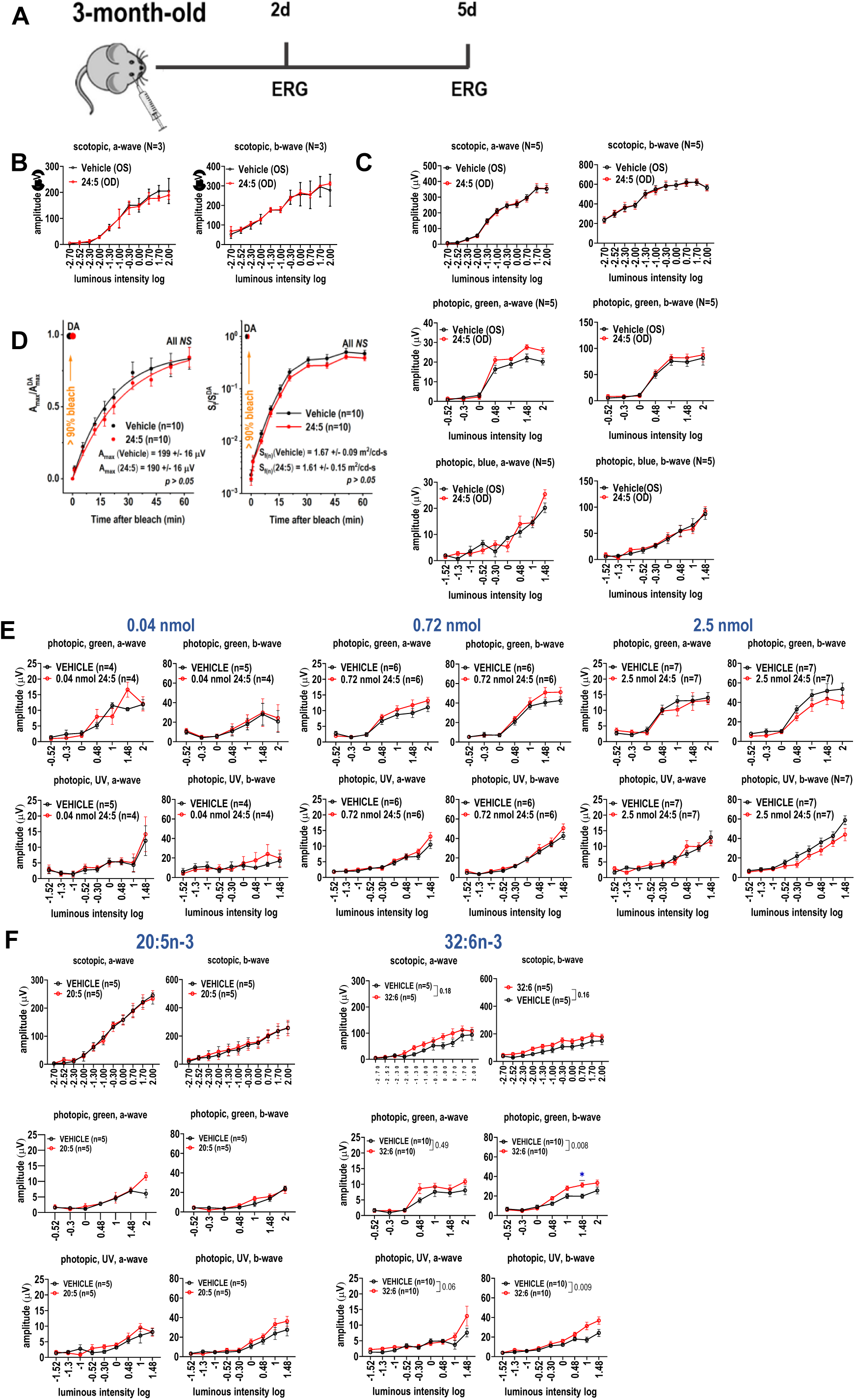
Lack of retinal toxicity or visual rescue in young mice following intravitreal supplementation of 24:5n-3 and optimal dosage of 24:5n-3, effect of different fatty acids on 18-month-old mice. **(A)** Schematic of intravitreal supplementation of 24:5n-3 in 3-month-old mice. **(B)** Electroretinogram (ERG) responses 2 days post-injection, **(C)** 5 days post-injection and **(D)** rod-mediated dark adaptation recovery in young animals remained unchanged after supplementation. **(E)** Photopic ERG responses 5 days post-injection in 18-month-old mice remained unchanged after supplementation with 0.04 nmol, 0.72 nmol and 2.5 nmol 24:5n-3, and (**F**) 0.36 nmol 20:5n-3 and 32:6n-3.

**Fig. S5.**
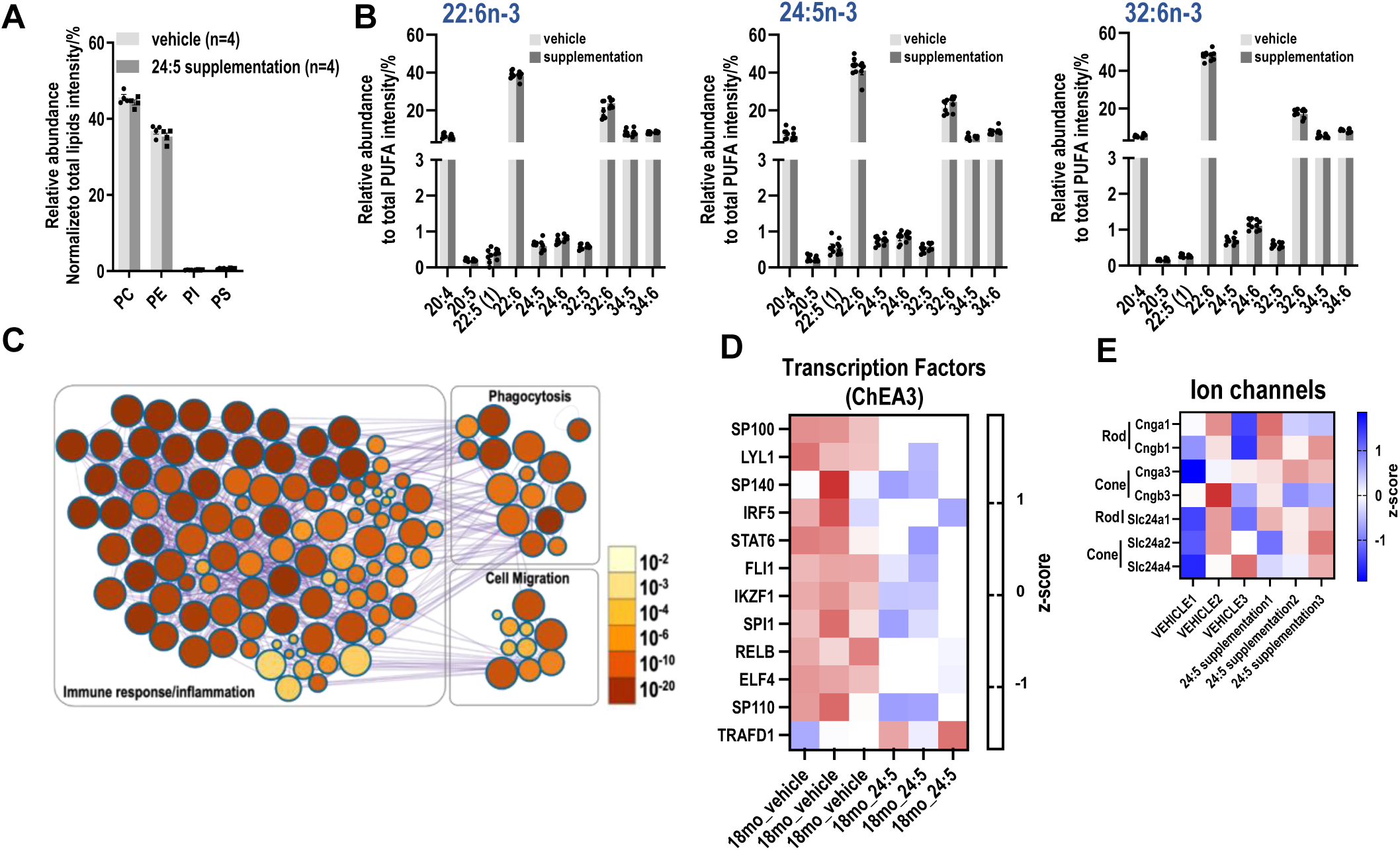
Lipid analysis in retina after lipid supplementation. (**A**) Major phospholipid classes remained unchanged in retinas from vehicle and 24:5n-3 supplemented retina. (**B**) Unchanged levels of PUFAs in retinas after 22:6n-3, 24:5n-3 and 32:6n-3 injection compared to vehicle injected (**C**) Metascape analysis of enriched clusters of pathways after 24:5n-3 supplementation colored by p-value. **(D)** Expression of TFs identified by ChEA3 after supplementation of 24:5n-3 **(E)** Heatmap expression levels of cone- and rod-specific ion channel genes with or without 24:5n-3 supplementation.

